# Robust detection of specific epistasis using rank statistics

**DOI:** 10.1101/2025.04.08.647864

**Authors:** Maryn O. Carlson, Bryan L. Andrews, Yuval B. Simons

## Abstract

The phenotypic effect of a mutation may depend on the genetic background in which it occurs, a phenomenon referred to as epistasis. One source of epistasis in proteins is direct interactions between residues in close physical proximity to one another. However, epistasis may also occur in the absence of specific interactions between amino acids if the genotype-to-phenotype map is nonlinear. Disentangling the contributions of these two phenomena—specific and global epistasis—from noisy, high-throughput mutagenesis experiments is highly non-trivial: the form of the nonlinearity is generally not known and model misspecification may lead to over- or underestimation of specific epistasis. In contrast to previous approaches, we do not attempt to model the fitness measurements directly. Rather, we begin with the observation that global epistasis, under the assumption of monotonicity, imposes strong constraints on the rank statistics of a combinatorial mutagenesis experiment. Namely, the rank-order of mutant phenotypes should be preserved across genetic backgrounds. We exploit this constraint to devise a simple semi-parametric method to detect specific epistasis in the presence of global epistasis and measurement noise. We apply this method to three high-throughput mutagenesis experiments, uncovering known protein contacts with similar or higher accuracy than existing, more complicated procedures. Moreover, the principles underlying our framework may suggest new ways of understanding the mechanisms which generate epistasis and their consequences for protein evolution.

## 1 Introduction

When two mutations are *epistatic*, the phenotypic effect of combining the two mutations cannot be predicted from their single effects alone. This non-independence of mutational effects is ubiquitous across biological systems and scales (Wolf et al., 2000; Phillips, 2008), impeding our ability to predict phenotypes from genotypes and the outcomes of sequence evolution (Papkou et al., 2023). How epistasis deforms the fitness landscape has been the subject of much theoretical work (Wright et al., 1932; Wolf et al., 2000; Weinreich et al., 2005, 2006). For example, positive epistasis among beneficial mutations may expedite the climbing of fitness peaks, while sign epistasis may impede the crossing of fitness valleys (Weinreich et al., 2005). Empirical characterization of fitness landscapes in various experimental settings has begun to test the relevance of these theoretical expectations to natural systems (e.g., Olson et al. 2014; Diss and Lehner 2018; Pokusaeva et al. 2019; Moulana et al. 2023; Westmann et al. 2024; Metzger et al. 2024).

The question of *measurement scale* has been central to the definition and detection of genetic interactions since R.A. Fisher introduced the notion of “epistacy” in the context of quantitative traits (Fisher, 1919; Cordell, 2002). For example, the analysis of genetic effects which combine mulitplicatively on an additive scale would result in the appearance of widespread interactions—a phenomenon Fisher referred to as “metrical bias” (Fisher et al., 1932). He supposed that metrical bias could often be removed by applying an appropriate, likely rank-preserving, transformation (Fisher et al., 1932; Mather, 1943; Crow, 1990; Phillips, 2008).

The question of the appropriate measurement scale continues to animate contemporary studies of epistasis, particularly in proteins. In proteins, information about epistasis comes from two primary sources: Multiple sequence alignments (MSAs) within protein families, and targeted mutagenesis of particular protein sequences. Analyses of MSAs have provided evidence for the evolutionary coupling of subsets of protein residues implicated in protein folding and function (Lockless and Ranganathan, 1999; Morcos et al., 2011; Salinas and Ranganathan, 2018; Wang et al., 2019). In contrast, targeted mutagenesis interrogates the local epistatic landscape more directly, by generating mutants within the neighborhood of a wildtype sequence(s) (e.g., Araya et al. 2012; Olson et al. 2014; Salinas and Ranganathan 2018; Poelwijk et al. 2019). Contemporary experiments, referred to as deep mutational scans (DMSs), assay the phenotypes of thousands to millions of mutants simultaneously with high throughput sequencing-based methods (Fowler and Fields, 2014; Kinney and McCandlish, 2019).

In DMSs assaying the combined fitness effects of two or more mutations (Fig. 1a-b), observed epistasis is often categorized into two types: *specific epistasis* (SE), where the effect of a mutation at one position depends on the identity of the amino acid at another via a direct interaction; and *global epistasis* (GE)—the analog of Fisher’s metrical bias—where apparent interactions between mutations emerge from the presence of nonlinearities in the genotype-to-phenotype map (Sailer and Harms, 2017a; Starr and Thornton, 2016; Husain and Murugan, 2020). The former, SE, is most often associated with amino acids in close proximity in the protein structure (Fig. 1c; Rollins et al. 2019; Schmiedel and Lehner 2019), while GE may be attributed to many causes, including a thermodynamic equilibrium between conformational states or detection limits imposed by an experimental assay (Fig. 1d, e.g., Sailer and Harms 2017a; Husain and Murugan 2020; Faure and Lehner 2024 and also see Lehner 2011; Johnson et al. 2023). Another body of work suggests that GE can arise from widespread SE (Kryazhimskiy et al., 2014; Lyons et al., 2020; Reddy and Desai, 2021), a scenario which we return to in Section 4. The extent to which SE and GE contribute to the epistasis observed in a given experiment (Fig. 1e) and, in turn, shape protein fitness landscapes is an open question (see Dupic et al. 2024 in response to Park et al. 2022; Faure et al. 2024) with consequences for how proteins evolve and function (Sailer and Harms, 2017b; Sato and Kaneko, 2020; Husain and Murugan, 2020).

**Figure 1.**
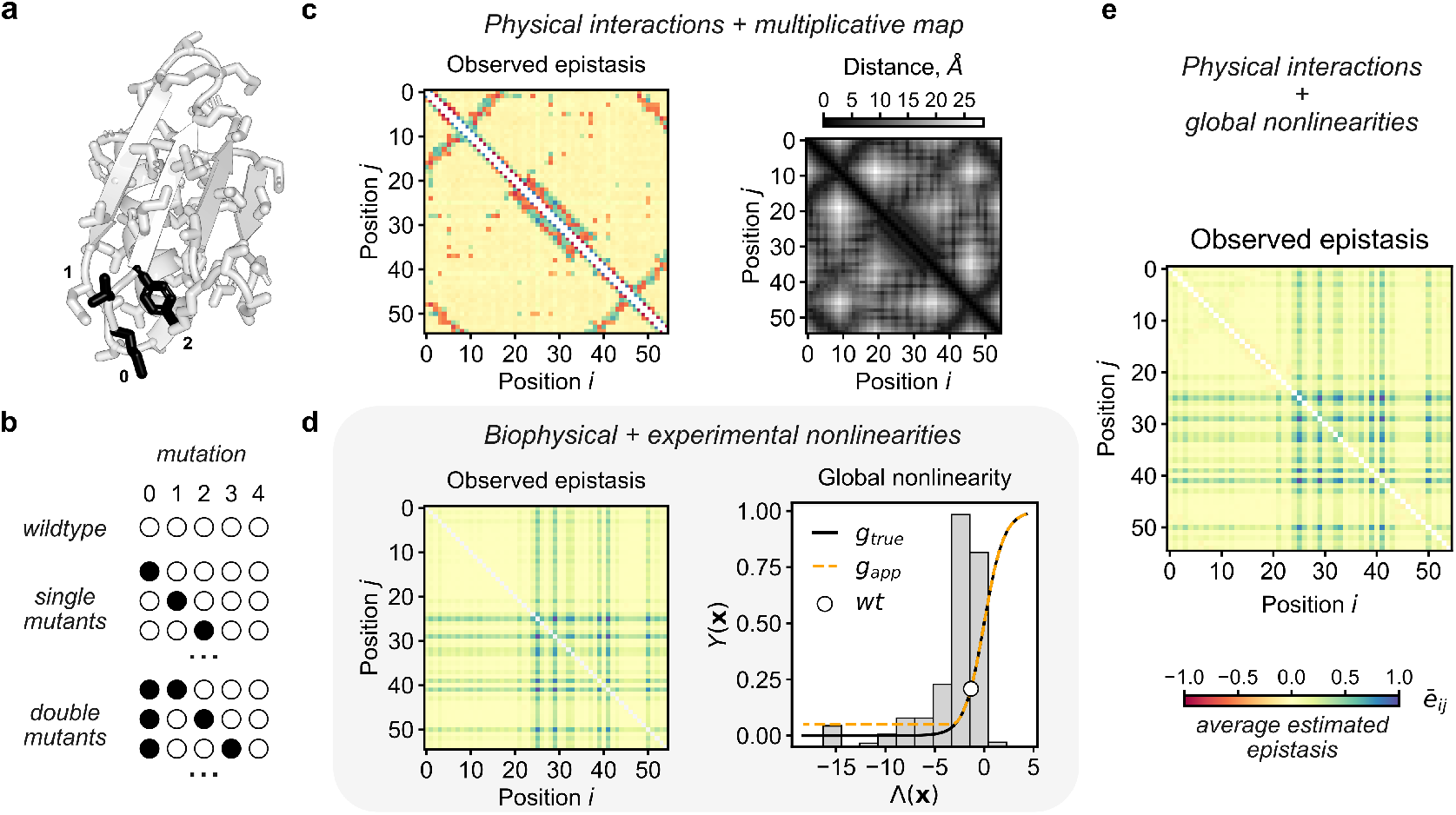
Potential sources of observed epistasis in proteins. **(a)** A linear sequence of amino acids folds into a 3-dimensional structure, bringing the amino acids at particular positions into direct contact with one another (PDB: 2J52, Wilton et al. 2008). **(b)** In a deep mutational scan, mutations (black circles) are introduced into a wildtype background (white circles). We will focus on DMSs that generate all or most of the single and double mutants of a given wildtype sequence. **(c)** If the genotype-to-phenotype map is multiplicative, we expect the detected epistasis, *ē*_*ij*_ (*left*), to reflect specific epistasis induced by the contact map (*right*, derived from **a**). Epistatic effects are computed with respect to a multiplicative fitness map, averaged over all pairs of amino acids for each position pair, and normalized by the maximum |*ē*_*ij*_ | for visualization. **(d)** Nonlinearities in the genotype-to-phenotype (**x** to Λ to *Y*) map of biological (*right, g*_true_, solid black line) and/or experimental origin (*g*_app_, dashed orange line), may introduce higher-order epistasis (*left*), even in the absence of specific interactions. Histogram of the simulated single mutant Λ values in gray (*right*). **(e)** Observed epistasis when both physical interactions and global nonlinearities determine the genotype-to-phenotype map.

Formally, under a model of GE, each single mutation *i* has an independent effect *λ*_*i*_ on a latent additive trait Λ (Fig. 1d). For example, Λ may correspond to the energy associated with protein folding or ligand binding. In addition, a sparse set of mutation pairs *i* and *j* may also exhibit second-order effects, *λ*_*ij*_, on the additive trait (Eq. 1a). For exposition, let **x** ∈ {0, 1} ^*L*^ be a binary, *L*-length protein sequence. In practice, mutations often occupy a larger state space (e.g.,the 20 amino acids).

The estimated phenotype associated with **x**, *Ŷ* (**x**), is a nonlinear *monotonic* function *g* of Λ(**x**),

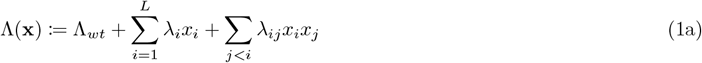

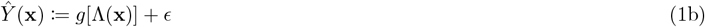

where Λ_*wt*_ is the value of Λ for the wild-type sequence, and *ϵ* is the measurement error with a potentially unknown distribution (see e.g., Otwinowski et al. 2018). In the event that *g* is nonlinear, estimation of the epistatic coefficients under the assumption of an additive model—equivalent to assuming that *g* is a linear function—may lead to spurious inference of many non-zero higher order coefficients. More broadly, misspecification of the form of *g* is apt to distort features of the genotype-to-phenotype map, over- or under-representing the importance of higher-order interactions (Sailer and Harms, 2017a; Otwinowski et al., 2018; Otwinowski, 2018).

For exposition, consider the example in Fig. 1, where the genotype-to-phenotype map coheres with the definition of GE given in Eq. 1, and a sparse set of interactions, *λ*_*ij*_ ≠ 0, corresponds to physical contacts within the protein (Fig. 1c). If mutations combine multiplicatively, the epistatic coefficients estimated under the assumption of a multiplicative model accurately reproduce the contact matrix (Fig. 1c). The presence of a nonlinearity induces pervasive epistasis in the absence of SE (Fig. 1d) and obscures SE when both are present (Fig. 1e). Accurately estimating the form of the nonlinearity *g* would, in principle, allow for a more parsimonious representation of the genotype-to-phenotype map (Otwinowski et al., 2018).

Thus motivated, researchers have developed several methods to account for GE when estimating epistatic coefficients or predicting mutant phenotypes. Several of these methods rely on strong assumptions about the form of the nonlinearity: for example, assuming that *g* is logistic (Park et al., 2022), follows from a thermodynamic model (Diss and Lehner, 2018; Otwinowski, 2018), or is otherwise of a pre-specified form (Sailer and Harms, 2017a; Pokusaeva et al., 2019; Faure and Lehner, 2024). Although other procedures impose fewer modeling assumptions, they do not provide a principled hypothesis testing framework for identifying epistasis (Schmiedel and Lehner, 2019), or they involve potentially cumbersome fitting procedures (Otwinowski et al., 2018; Tonner et al., 2022; Faure and Lehner, 2024).

Indeed, developing a hypothesis testing framework in the presence of GE is highly nontrivial. Firstly, fitness measurements are often derived from noisy, sequencing-based assays with systematic variation in precision across the measurement range. This heteroskedasticity implies that statistical power to detect SE in a well-calibrated statistical test should similarly vary across the measurement range. Secondly, if not properly accounted for, uncertainty in the estimation of *g* may lead to over- or underestimation of the prevalence of SE.

In contrast to all previous approaches for detecting SE in the presence of GE, we do not attempt to explicitly estimate the form of the nonlinearity, nor to model the fitness measurements directly. We do not even assume that *g* is nonlinear. Rather, our work begins with the observation that GE imposes strong constraints on the rank statistics of a DMS. Namely, if the latent space is uni-dimensional, which we will assume going forward (though see Section 3.3 and Section S2.4.1), and if the nonlinearity *g* is monotonic and strictly increasing (or decreasing), then *g* is also order preserving in the absence of measurement noise. In other words, when *g* is monotonic, the ordering of mutations is shared across genetic backgrounds. This observation similarly motivated Brookes et al. (2024) to introduce a rank-based loss function for phenotypic prediction in the context of GE. Our work further exploits the (assumed) monotonicity of GE to detect SE, which disrupts the ordering of mutations.

We first demonstrate that rank statistics are a natural framework for the analysis of combinatorial data sets, a notion which has only recently been developed in the literature (Crona et al., 2017; Lienkaemper et al., 2018) and applied to protein DMSs (Brookes et al., 2024). We then define a rank-based measure of SE and use it develop a semi-parametric test for deviations of mutation pairs from GE that accounts for heteroskedasticity, which may arise due to variation in the shape of *g* or scale of the measurement noise (or both) across the measurement range. Our procedure—referred to as Resample and Reorder or R&R—requires minimal pre-processing of the data beyond generating variant read counts, is invariant under monotonic transformations of the data, and is agnostic to the form of the non-linearity beyond monotonicity. We apply R&R to simulated and empirical DMSs of proteins, demonstrating its ability to recover true epistatic effects and physical contacts, respectively. Finally, we explore the consequences of misspecification of the GE model on the results of our inference procedure. In particular, we consider a scenario where there are two, rather than one, latent additive traits. While we motivate R&R using DMSs of proteins, the method generalizes almost immediately to other types of combinatorial data sets.

## 2 Modeling framework

### 2.1 Rank statistics as a natural framework for global epistasis

We begin with the observation that GE imposes strong constraints on the rank statistics of a combinatorial mutagenesis experiment, such as a DMS: the fact that the nonlinearity *g* is monotonic implies that the rank-order of mutations should be preserved regardless of the background in which the mutations occur. As a consequence, in the absence of measurement noise and SE, the Spearman’s correlation between mutant phenotypes measured in distinct backgrounds *i* and *j*, 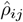, is equal to one. Measurement noise, however, may result in Spearman’s correlations that deviate substantially from one, even in the absence of SE (Fig. 2b).

**Figure 2.**
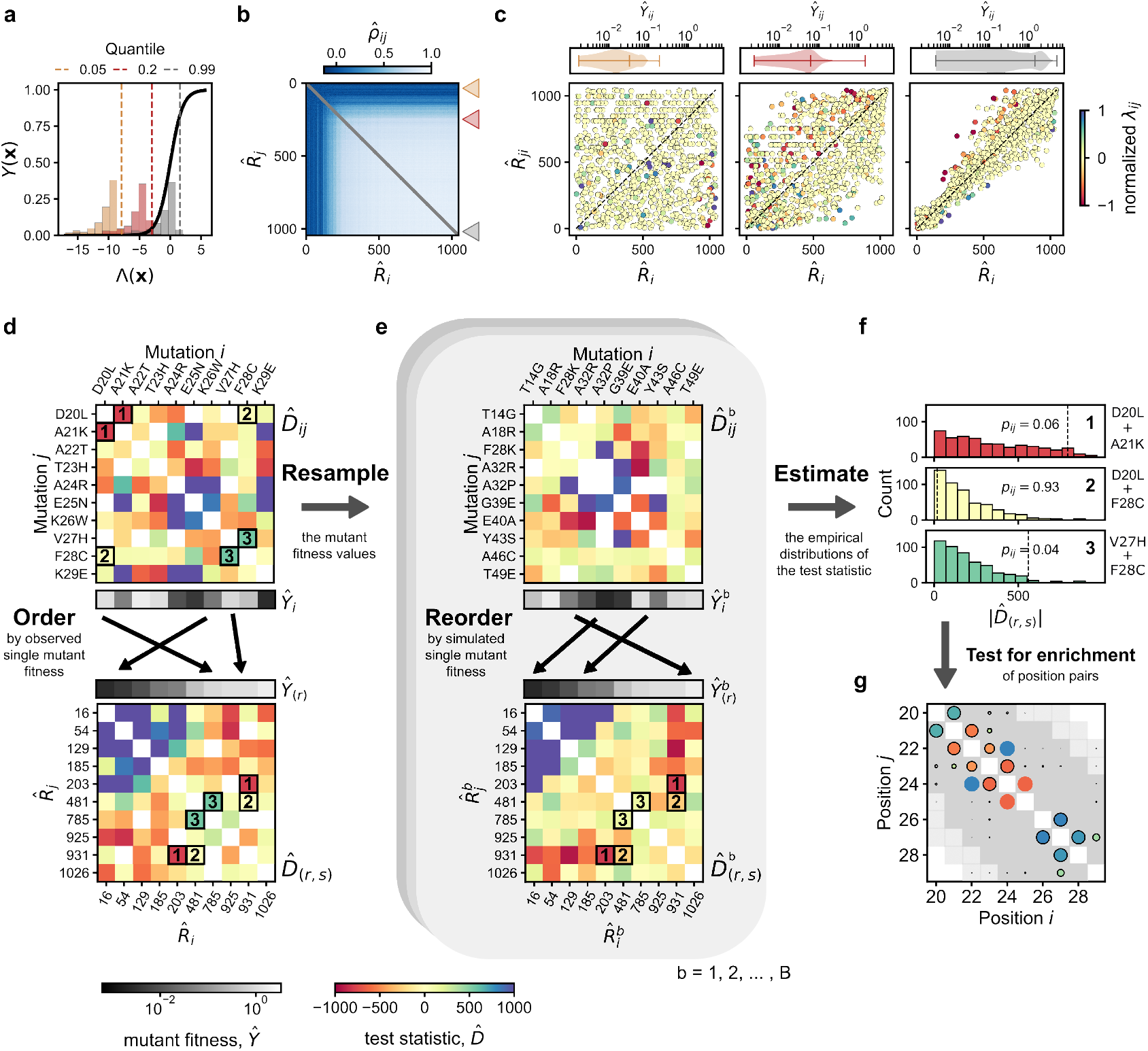
Rank statistics as a natural framework for detecting specific epistasis in the presence of global epistasis. **(a)** Probability of binding, *Y*, as a function of the latent trait Λ (solid black line). Three focal mutations, in brown, red, and gray, with distinct Λ values (dashed vertical lines) from a DMS simulated under the assumption of a two-state thermodynamic model. Histograms correspond to the Λ values of double mutations containing the focal mutations. **(b)** Spearman’s correlation matrix for all pairs of mutations *i* and *j*, computed from the estimated double mutant phenotypes *Ŷ*_*ij*_, and ordered by single mutant ranks 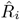. **(c)** Distributions of *Ŷ*_*ij*_ for each of the three focal backgrounds (*top*). Double mutant ranks 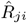 for each of the three backgrounds as a function of the single mutant ranks 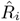 (*bottom*). Each point is colored by its true specific epistatic effect, *λ*_*ij*_. **(d)** Observed values of 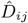 for a subset of mutations at nearby positions (positions 20-29) in position order (*top*). Mutations are reordered (*bottom*) by their single mutant fitness values *Ŷ*_*i*_, denoted *Ŷ*_(*r*)_ (gray scale heatmap). Three focal rank pairs, 1 − 3, are outlined with solid black squares. **(e)** New values of *Ŷ*_*i*_ and *Ŷ*_*ij*_ are generated under the assumption of a particular error model (*top*), shuffling which mutations are associated with a given rank in a given bootstrapped replicate *b* (*bottom*). **(f)** Empirical distributions of 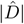 plotted for the three focal pairs denoted in **d. (g)** Position pairs enriched for significant interactions colored by average sign and shown with respect to physical contacts (≤ 5Å, dark gray) and nearby positions (≤ 8Å, light gray). The size of the point is proportional to the −*log*_10_ *p*-value of enrichment.

We focus on a class of DMSs in which the phenotypes of single and double mutants have been estimated. Consider the ordering of two mutants *m* and *n* in the background of mutant *i*. We denote their true fitness values as *Y*_*im*_ and *Y*_*in*_. If the difference between *Y*_*im*_ and *Y*_*in*_ is small relative to the magnitude of the measurement noise, *σ*_*ϵ*_, then each of the possible orderings of their *estimated* fitness values, *Ŷ*_*im*_ > *Ŷ*_*in*_ and *Ŷ*_*im*_ *< Ŷ*_*in*_, is approximately equally likely. Such small differences arise when (1) differences in the mutations’ effects on the latent trait, *λ*_*m*_ and *λ*_*n*_, are small; (2) the slope of the nonlinearity *g* in the neighborhood of the background *i* is small; or, (3) *σ*_*ϵ*_ is large. More succinctly,

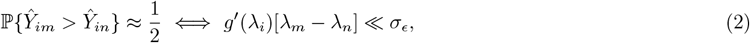

where, for the sake of exposition, we have assumed that both *λ*_*m*_ and *λ*_*n*_ are small (Section S2.6.2). Thus, in general, in the absence of SE, deviations of 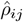 from one will be due to the mis-ordering of mutants with small differences in their latent effects relative to the measurement noise. In addition, variation in the slope of the nonlinearity and the magnitude of noise across the measurement range may induce systematic variation in 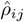 as a function of single mutant fitness.

To illustrate how the presence of a nonlinearity and noise introduce variation in the 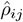 values among mutants, we simulate a DMS under the assumption of a two-state thermodynamic model where single mutant effects on binding are specified by their estimated values from Otwinowski (2018) (Fig. 2a). An important feature of this model is saturation at both very small and large values of Λ: in the background of a very deleterious mutation, additional deleterious mutations will not further reduce fitness. Similarly, in the background of a very fit mutation, a beneficial mutation will not confer an additional fitness advantage—a form of diminishing returns epistasis.

At these two extremes, differences in fitness among mutants will be small relative to the magnitude of measurement noise, implying that the double mutant phenotypes will be approximately uniformly ordered. This implies that for a very deleterious (or beneficial) mutant *i*, 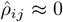 for all mutations *j* ≠ *i*. As a consequence, ordering the Spearman’s correlation matrix by single mutant rank reveals a characteristic structure, with row maximum (and mean) systematically increasing with single mutant rank (Fig. 2b and see Section S2.6). Similarly, the estimated rank of a mutation *m* in the background of a very deleterious mutant *i*, 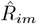, will be uncorrelated with its estimated single mutant rank, 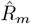 (Fig. 2c). In contrast, mutations in the background of a typical mutation will be well-ordered, with small deviations due to measurement noise and large deviations due to SE (Fig. 2c). We will exploit the latter feature to detect SE.

### 2.2 Detecting specific epistasis

We formally define SE as the presence of a non-zero interaction between two mutations *i* and *m*, with respect to the latent additive trait Λ, i.e., *λ*_*im*_ ≠ 0 (Eq. 1). If the magnitude of *λ*_*im*_ is large enough, it will lead to deviations from the expected ordering under a model of GE. For exposition, consider again the ordering of the phenotypes of two mutants *m* and *n* in the background of mutant *i*. Suppose, without loss of generality, that *λ*_*n*_ > *λ*_*m*_ and, in accordance, the estimated fitness of single mutant *n* exceeds that of mutant *m*, i.e., *Ŷ*_*n*_ > *Ŷ*_*m*_. If mutations *i* and *m* are epistatic, and *λ*_*im*_ > 0, the probability that *Ŷ*_*im*_ is greater than *Ŷ*_*in*_ can be approximated by,

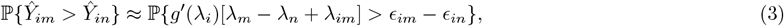

when *λ*_*m*_, *λ*_*n*_, and *λ*_*im*_ are small (and *λ*_*in*_ = 0); and, *ϵ*_*im*_ and *ϵ*_*in*_ are the measurement errors of the double mutants (Section S2.6.6). Therefore, when the epistatic effect *λ*_*im*_ exceeds a threshold set by the measurement noise, the local curvature of the nonlinearity, and the difference in first order fitness effects, i.e.,

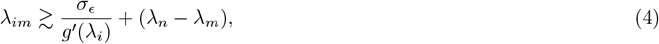

SE will likely result in a change in the ordering of the double mutants with respect to that of the single mutants: *Ŷ*_*im*_ > *Ŷ*_*in*_ while *Ŷ*_*m*_ *< Ŷ*_*n*_ (Section S2.6.7). Moreover, for any *n* such that Eq. 4 holds, *Ŷ*_*im*_ > *Ŷ*_*in*_ is a likely outcome. Thus, a natural summary of the magnitude of SE in the context of rank-statistics is the difference between the rank of the double mutant *im* compared to the rank of the single mutant *m*. Below, we propose a simple semi-parametric test of SE based on this precept that makes no assumptions about *g* beyond monotonicity.

#### 2.2.1 A rank-based test statistic

The aim of our inference procedure is to assess whether or not a given pair of mutations exhibits SE while allowing for the presence of a global nonlinearity (Eq. 1). Namely, we want to conduct a hypothesis test, where the null hypothesis is given by:

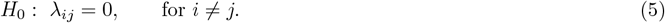

Under the null hypothesis and the assumption of GE, and in the absence of measurement noise, the rank of mutant *j* in the background of mutant *i*, denoted as *R*_*ij*_, is equal to the rank of mutant *j* in the wildtype background, *R*_*j*_. More concisely, *R*_*ij*_ = *R*_*j*_. Thus, a natural, rank-based estimate of the epistatic effect between mutants *i* and *j, λ*_*ij*_, is the difference in the estimated double and single mutant ranks. We define the test statistic 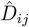 as the sum of the rank differences between the double and single mutants for mutants *i* and *j*, respectively,

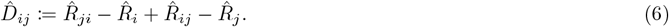

When calculating 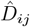 in practice, we account for missing data by transforming the ranks onto the same scale (Section 5.2.1 and see Section S1.2.1). In Fig. 2d, we compute 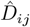 for a subset of mutations with varying fitness values.

A deviation of 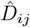 from zero would naïvely provide evidence for SE between mutants *i* and *j*. The challenge in testing for significant deviations of 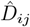 from zero (and thus *λ*_*ij*_ from zero) is that in the presence of measurement noise and GE, the variances of the constituent quantities may vary systematically across the measurement range (Fig. 2c). In addition, even in the absence of SE, the expected value of 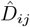 is not necessarily equal to zero. For example, the value of the test statistic for the two most deleterious mutants will be non-negative, as ranks are bounded below by zero. Thus, correct calibration of the proposed hypothesis test requires estimation of the distribution of 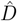 for each pair of ranks, 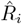 and 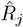, in the presence of measurement noise.

#### 2.2.2 Resample and reorder, a bootstrapping approach

To estimate the null distributions of the test statistics for each pair of single mutant ranks, we take a simple, yet novel bootstrapping approach. The bootstrapping procedure requires specification of an error model—the semi-parametric component of R&R. The choice of error model, however, is not conceptually fundamental to our approach and necessarily depends on the application. When fitness estimates are defined by the ratio of sequencing reads after and before selection—as in the DMSs considered in this study—a Poisson error model is a natural choice (Faure et al. 2020 and see Section 5.2.2).

In each bootstrap replicate *b* = 1, …, *B*, we generate a synthetic data set encompassing fitness estimates for all observed single and double mutants (Fig. 2e, *top*). The mutants are then ranked according to their fitness estimates, generating a new set of single mutant ranks, 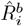, for all mutants *i* = 1, …, *M*, and a new matrix of double mutant ranks, 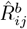 for *j* ≠ *i*. The test statistic is then computed for every pair of mutations in each replicate, yielding 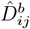 values for all possible pairs of mutations *i* and *j* (Fig. 2e, *bottom*).

The key feature of our procedure is that the mutants with given ranks *r* and *s* will vary across simulations (Fig. 2e). This shuffling allows us to estimate the distribution of 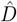 as a function of the *ranks* of the constituent mutations, rather than their identities, ultimately yielding empirical distributions of the absolute value of the test statistic, 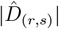, for each pair of mutant ranks *r* and *s* (Fig. 2f). The *p*-value for a given pair of mutations with observed ranks 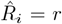 and 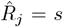 is then computed with reference to the empirical distribution of the test statistic 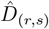 (Fig. 2f and Eq. 10). To identify SE at the level of position pairs, and to overcome the multiple testing burden, each pair of positions is tested for enrichment of *p*-values below a fixed threshold *α* among all observed amino acid combinations at the two positions (Fig. 2g and see Section 5.2.5).

## 3 Evaluating R&R on simulated and empirical datasets

### 3.1 Resample and reorder identifies true epistatic effects in simulations

To evaluate R&R, we again simulate a DMS where the global nonlinearity is specified by a two-state thermodynamic model (Section 5.3.1). The simulations are motivated by the DMS of Olson et al. (2014) in which the authors generated all single and most double mutants of the IgG-binding domain of protein G (GB1), and a subsequent analysis of these data by Otwinowski (2018).

We first consider an ideal scenario in which (1) all single and all double mutants are represented at equal frequencies in the initial library, respectively, and (2) the effects *λ*_*i*_ on the latent trait Λ are uniformly distributed (Fig. 3a, left). The first simulation condition, equal cell counts, implies that the phenotypic measurements, *Ŷ*_*i*_ and *Ŷ*_*ij*_, are independent and identically distributed (*iid*) random variables when conditioned on the latent trait values, Λ_*i*_ and Λ_*ij*_, respectively. The second simulation condition, uniform sampling of the Λ_*i*_ values, ensures that variance in statistical power across the measurement range is not due to variation in sampling of the latent space. To introduce SE, we sample the epistatic coefficients *λ*_*ij*_ of position pairs within 5Å from a distribution with an expected value that decays as a function of the physical distance between positions. All other *λ*_*ij*_ values are set to zero (Section 5.3.2). To simulate the “observed” fitness estimates for each single and double mutant, pre- and post-selection read counts are sampled from Poisson distributions parameterized by each mutant’s initial cell count and true fitness (Fig. 3a). To apply R&R, we generate *B* = 1000 bootstrap samples, where in each bootstrap replicate, the pre- and post-selection read counts for all mutants are sampled from Poisson distributions with means given by their observed counts (Section 5.2.2). For more detailed simulation procedures, see Section 5.3.

**Figure 3.**
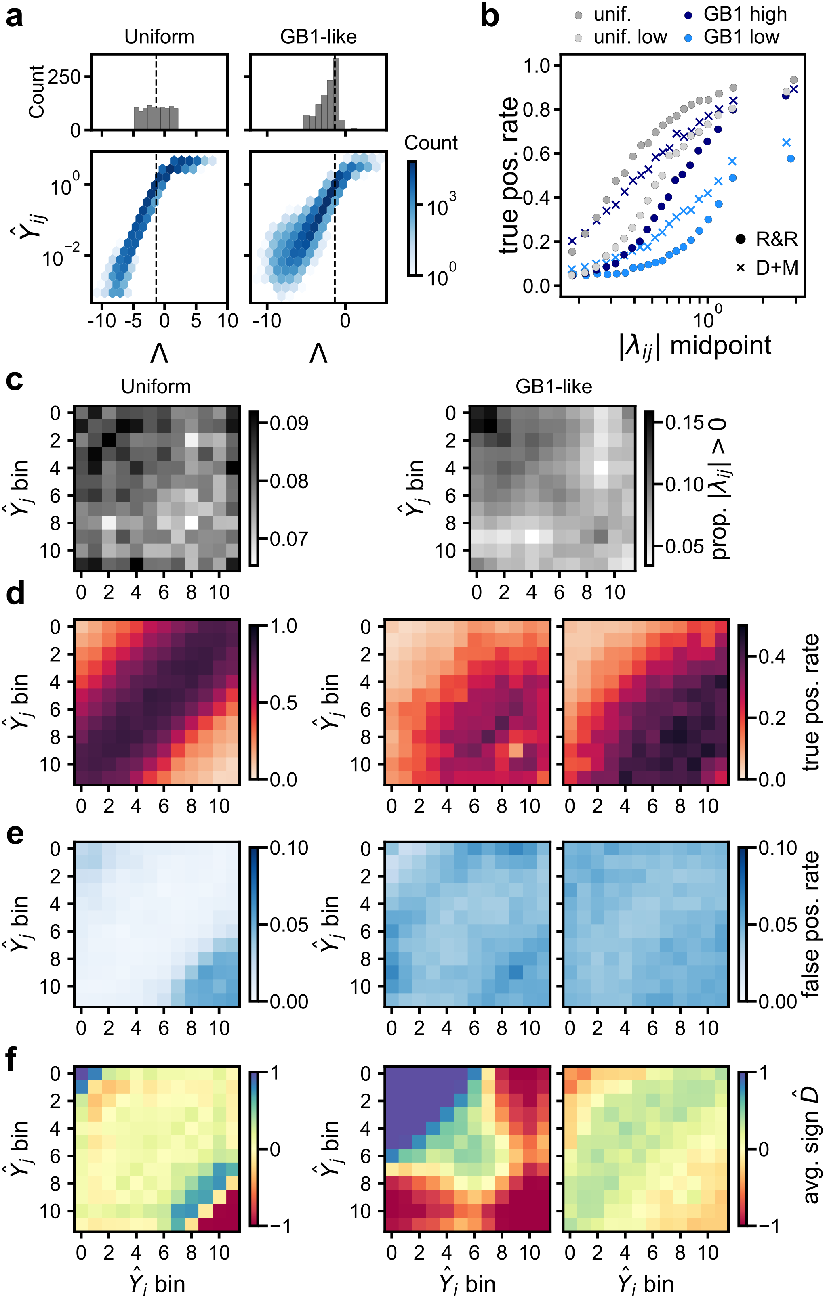
Resample and reorder accurately identifies epistatic effects in simulations. Estimated single and double mutant phenotypes were simulated under the assumption of a two-state thermodynamic model with epistatic effects sampled in accordance with the contact map. **(a)** Histograms of the single mutant latent trait values Λ_*i*_ (*top*), and double mutant phenotypes as a function of their Λ_*ij*_ values (*bottom*) in the *uniform* (*left*) and *GB*1-like simulations (*right*). The wildtype latent trait value, Λ_*wt*_, denoted by a black dashed line. **(b)** True positive rate (*tpr*) as a function of the magnitude of *λ*_*ij*_ for the *uniform* (gray), *GB*1-high (blue), and *GB*1-low simulations (orange) analyzed with R&R (circles) and `D+M` (x’s). **(c)** The proportion of non-zero interactions, *λ*_*ij* ≠_ 0 among positions within evenly spaced single mutant fitness *Ŷ*_*i*_ bins for the *uniform* (*left*) and *GB1-like low* (*right*) simulations. **(d)** True positive rate for each (*Ŷ*_*i*_, *Ŷ*_*j*_) bin pair for R&R analysis of the *uniform* (*left*) and *GB1-like low* (*middle*) simulations, and `D+M` analysis of the latter (*right*). **(e)** False positive rates for the two simulation scenarios and methods, as in **d. (f)** Average sign of significant test statistics for the simulations and methods, as in **d**.

As expected, the true positive rate increases as a function of the magnitude of the epistatic effect *λ*_*ij*_ (Fig. 3b). However, statistical power to detect significant associations is not uniformly distributed across the measurement range, despite the uniform distribution of true epistatic effects. Power is lowest for combinations of poorly or highly ranked single mutations (Fig. 3c-d). Two factors likely contribute to the systematic reduction in power for these extreme pairs. First, in the two-state model, the slope of the nonlinearity approaches zero for both deleterious and beneficial mutations. As a consequence, the variance of the test statistic is large for extreme pairs. Second, even in the absence of saturation, power to detect beneficial or deleterious interactions is asymmetric. A deleterious interaction (*λ*_*ij*_ *<* 0) will only minimally reduce the fitness of a double mutant composed of two very deleterious single mutations (and likewise for a beneficial interaction, *λ*_*ij*_ > 0, between two beneficial single mutations), resulting in systematic biases in the inferred *sign* of epistasis (Fig. 3f). The extreme pairs also contribute disproportionately to the false positive rate, presumably due to greater variance of the test statistic (Fig. 3e).

Next, we introduce variation in measurement precision by varying the initial cell counts, as well as a small probability of non-specific binding—both characteristic of empirical DMSs (Section 5.3). In addition, we set the single mutant effects on energy *λ*_*i*_ to their estimates from Otwinowski (2018) with some modifications (Fig. 3a, right and see Section 5.3.2). In Section S2.1, we consider each of these simulation amendments in isolation. Under this specification of *λ*_*i*_, more double mutants exhibit fitness values at the so-called lower detection threshold, resulting in post-selection counts at or near zero (Fig. 3a). And, unlike in our prior simulations, SE is concentrated among pairs of deleterious mutants, as mutations at positions engaged in many physical contacts tend to reduce fitness (Fig. 3c).

When R&R is applied to this GB1-like DMS, we observe a wholesale reduction in power relative to the uniform simulations (Fig. 3b,d). Reductions in power are exacerbated when we reduce the initial cell counts by a factor of ten, further decreasing measurement precision—a simulation scenario we refer to as GB1-low in Fig. 3b. In the GB1-like simulations, we observe more pronounced asymmetry in the ability of R&R to detect positive and negative epistasis in different areas of the measurement range. In particular, R&R almost exclusively detects negative epistasis among pairs of deleterious and beneficial mutants (Fig. 3f; GB1-low and see Fig. S2d for GB1-high). This asymmetry likely arises due to saturation at the low-end of the measurement range (see Section S2.6.8 for a more detailed discussion). To mitigate the sign bias, we consider a two-sided hypothesis test in Section S1.5.

To further benchmark R&R, we compare its performance to that of an existing procedure which combines the fitness estimates of `DiMSum` (Faure et al., 2020) with a neural network-based GE inference framework, `MoCHI` (Faure and Lehner, 2024), referred to henceforth as `D+M`. `D+M` tests for SE by examining the residuals of the double mutant fitness estimates with respect to a fitted GE model: large residuals, normalized by the standard error of the double mutant fitness estimate, provide evidence for SE. We apply `D+M` to the GB1-like simulations, specifying the form of GE as a sum of sigmoid functions. (See Section S1.4 for more details.)

The *p*-values of R&R and `D+M` are highly correlated (Spearman’s *ρ* ≈ .75 for both high and low simulations), though they exist on vastly different scales; the *p*-values of R&R are constrained by the number of bootstrap samples (Eq. 10), while those of `D+M` can, in theory, span the unit interval. To provide a meaningful comparison between the methods, we fix the false positive rate across the two methods in each simulation scenario (see Eq. S4). Under this parameterization, `D+M` is better powered to detect weak SE relative to R&R, with less appreciable differences in power for the GB1-low simulations (Fig. 3b). R&Rs reduced power relative to `D+M` is not surprising given overdispersion in the bootstrap sample (see Sections S2.1 and S2.2). Both methods are relatively less powered to detect interactions among deleterious mutants (Fig. 3d and Fig. S2d).

While `D+M` is better powered to detect SE in the simulation scenarios considered, we emphasize that R&R can achieve comparable results—particularly when fitness measurements are less precise—without estimating the non-linearity and at a fraction of the computational cost.

### 3.2 Resample and reorder identifies protein contacts in empirical data sets

Next, we quantify the ability of R&R to identify protein contacts in two DMSs, for which previous studies support models of single-trait GE and a strong association between inferred SE and physical proximity (Diss and Lehner, 2018; Schmiedel and Lehner, 2019; Zarin and Lehner, 2024). In the first, Diss and Lehner (2018) conducted a DMS of two alpha-helical proteins, Fos and Jun, which form a heterodimeric transcription factor *in vivo* (Fig. 4a). The authors estimated the interaction strength of all single mutants and the majority of double *trans*-mutants—pairwise combinations of single mutations in each protein—in a high-throughput, sequencing-based assay. In the second, Zarin and Lehner (2024) estimated the binding affinities of almost all single and the majority of double *trans*-mutants of the third PDZ domain of PSD-95, PDZ3, for its 8-residue cognate ligand CRIPT (Fig. 4f). Due to batch effects in the PDZ3-CRIPT data set, we separately analyze the first and latter 50 amino acids of PDZ3, referred to henceforth as blocks 1 and 2 (Fig. S16). In addition, due to systematic biases in read coverage (see Figure S4E of Zarin and Lehner 2024), we exclude 14 positions in PDZ3 (see Section 5.1.3 for additional details). Both experiments were repeated three times, allowing us to examine the reproducibility of the results of R&R across replicates.

**Figure 4.**
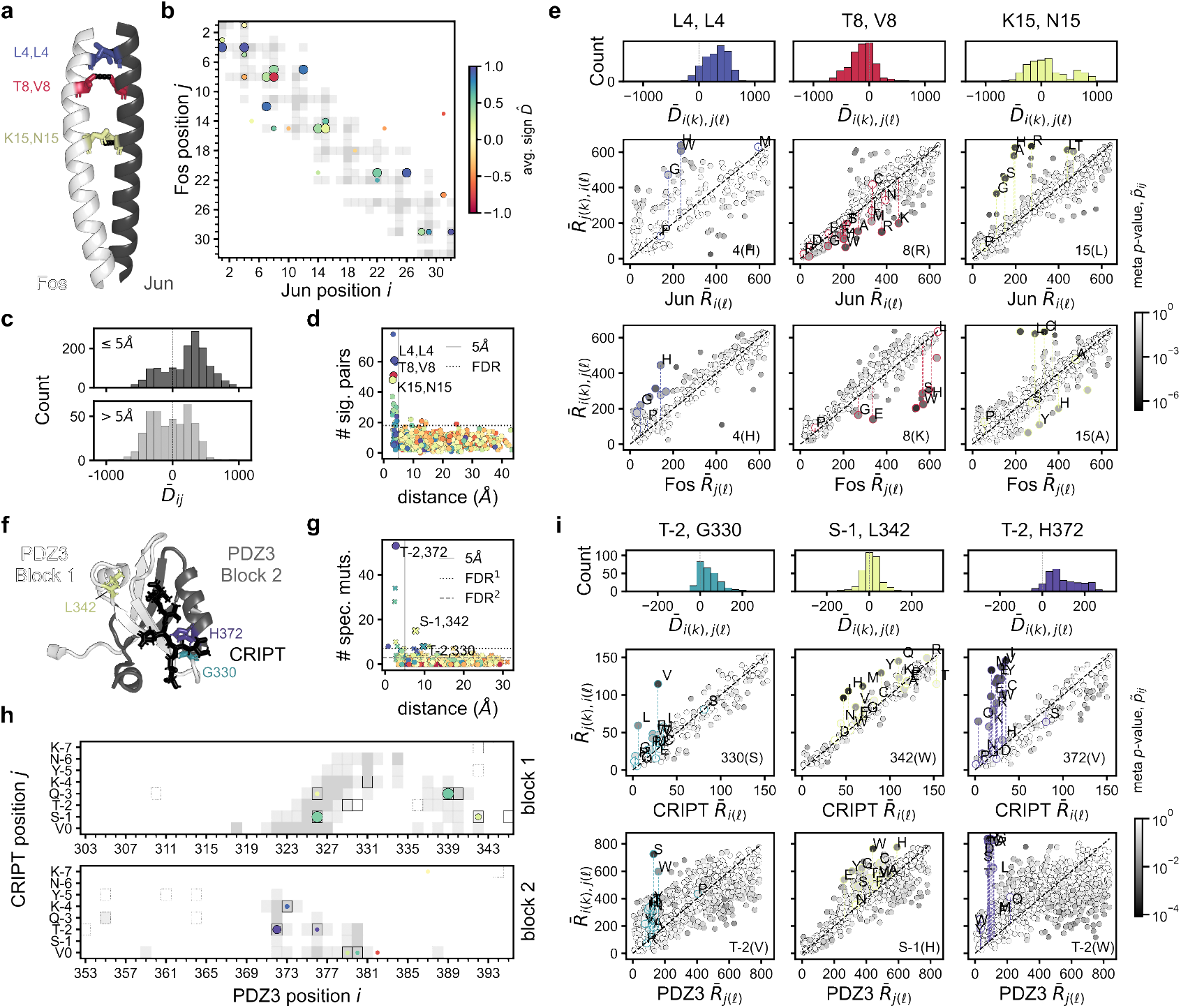
Resample and reorder identifies protein contacts in empirical data sets. **(a)** Crystal structures of the Fos-Jun complex (PDB: 1FOS, Glover and Harrison 1995). Three position pairs enriched for specific epistasis (SE) after correcting for multiple testing (*α*_*BH*_ ≤ .01) are highlighted and colored according to the average sign of the test statistic, 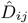. **(b)** All position pairs enriched for SE in the Fos-Jun dataset (*α*_*BH*_ = .1) are indicated by points, colored by the average sign of significant amino acid pairs, with the size of the point proportional to the −*log*_10_ enrichment *p*-value. Pairs significant at a false discovery rate of *α*_*BH*_ = .01 are additionally outlined in black. **(c)** Histograms of the average value of the test statistic, 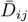, across replicates for significant amino acid pairs (*α* ≤ 10^−1^) among enriched protein contacts (*α*_*BH*_ = .1; *top*, dark gray) and positions at distances greater than 5Å (*bottom*, light gray). **(d)** The number of significant *p*-values (*α <* .1) associated with a given Fos-Jun position pair as a function of physical distance, with the dashed gray line indicating 5Åand the dotted black line denoting the significance threshold (*α*_*BH*_ = .1). **(e)** Each column corresponds to a different enriched position pair in Fos-Jun. Histograms of the corresponding 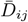 values, averaged over replicates (*top*), and average double mutant rank, 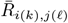, as a function of the single Jun (*middle*) and Fos (*bottom*) mutant ranks. The shading of each point denotes the meta *p*-value, 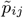. Points corresponding to the focal pairs are outlined in color and labeled with the associated amino acid. **(f)** Crystal structure of the PDZ3-CRIPT complex (PDB: 5HEB, Raman et al. 2016). Three significant position pairs are highlighted and colored according to the average sign, as in **a. (g)** The same plot as in **d**, except the number of *positive* interactions is shown for each PDZ3-CRIPT position pair, in keeping with (Zarin and Lehner, 2024). Block 1 and block 2 positions are represented by x’s and points, respectively. **(h)** All position pairs enriched for SE in PDZ3-CRIPT after multiple testing correction indicated by points as in **b**. In addition, position pairs enriched for *positive* SE at false discovery rates of *α*_*BH*_ = .1, .01 are denoted by dotted and solid black boxes, respectively. **(i)** The same plots as in **e** for select PDZ3-CRIPT position pairs.

We implement R&R with *B* = 1000 bootstrap replicates under the Poisson model (Section 5.2.2). In both data sets, we find that the *p*-values are approximately uniformly distributed in each replicate, with a more pronounced depletion of small *p*-values in the PDZ3-CRIPT dataset (Figs. S3 and S17), a phenomenon observed in simulations (Fig. S1). In addition, as in simulations, we observe variation in statistical power and the sign of detected SE—where pseudo-true positives are defined as positions pairs whose wildtype residues are within 5Å and whose *p*-values are below a fixed *α*—as a function of the single mutant ranks (Fig. 5a). In particular, R&R has lower pseudo-true positive rates among deleterious mutant pairs, which comprise the majority of protein contacts in both data sets (Fig. 5). Still, the total number of significant interactions is highest among deleterious mutant pairs in Fos-Jun (Fig. S10).

**Figure 5.**
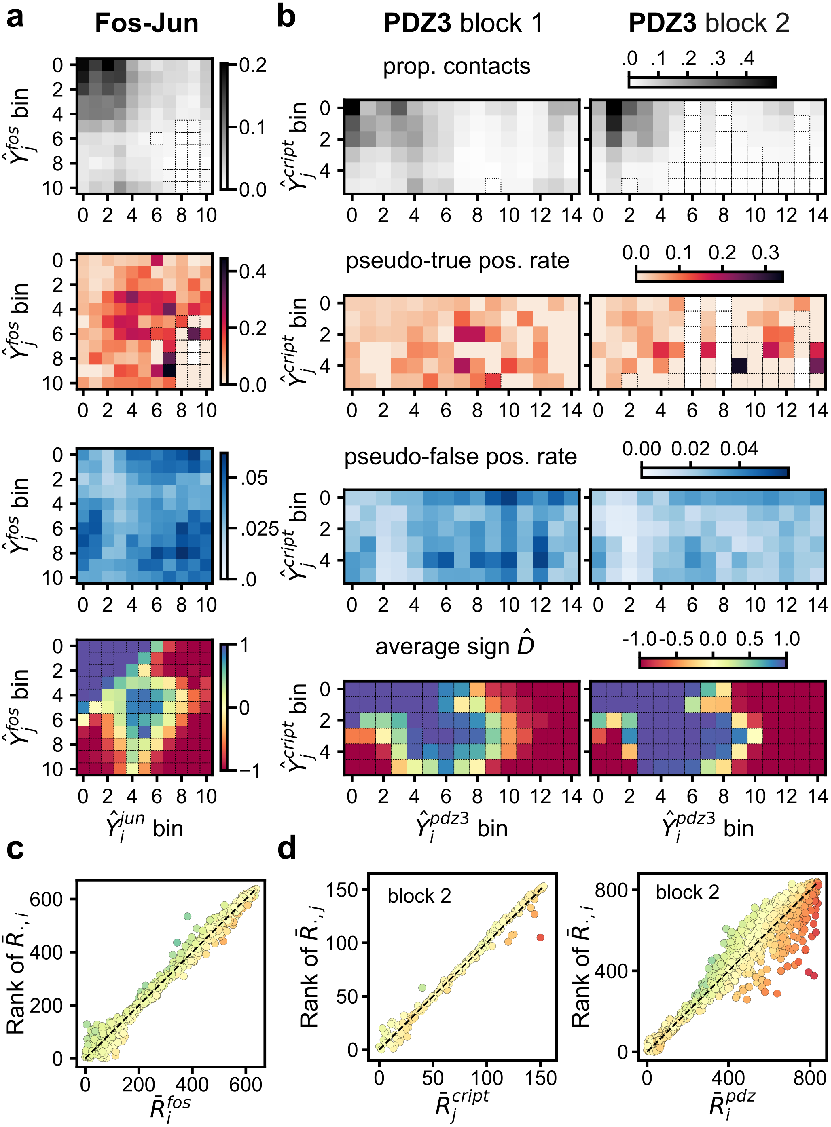
Evaluating R&R on empirical datasets. **(a)** A square in each heatmap represents a quantity averaged over all mutants in a pair of Fos and Jun single mutant phenotype, *Ŷ*_*i*_, bins. From the top, the proportion of contacts, the pseudo-true positive rate, the pseudo-false positive rate, and the average sign 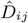 of significant amino acid pairs, all computed for *α* = 10^−1^. Dotted outlines indicate that the value was computed from fewer than five mutant pairs. **(b)** The same quantities computed for blocks 1 (left) and block 2 (right) of PDZ3 and CRIPT, with *α* = .043 and .028, respectively. **(c)** Average rank across mutational background plotted as a function of single mutant ranks Fos single mutants, averaged over replicates, with points colored by the average sign of corresponding significant 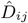 values, with color scale of **a-b. (d)** The same as **c** for CRIPT (*left*) and PDZ3 (*right*) single mutants in block 2.

Thus far, we have analyzed the *p*-values of position, amino-acid pairs. To identify interactions between positions, we follow previous work (Rollins et al., 2019; Schmiedel and Lehner, 2019; Zarin and Lehner, 2024) and test whether specific position pairs are enriched for SE, defined as mutant amino-acid pairs with *p*-values below a threshold *α* (Section 5.2.5).

#### Fos-Jun

Consistent with prior studies (Schmiedel and Lehner, 2019), enriched position pairs are sparse and predominantly constrained to protein contacts, defined as amino acids within 5Å in the crystal structure (Fig. 4b,d) with R&R achieving comparable or higher contact prediction accuracy relative to previous analyses (Diss and Lehner, 2018; Rollins et al., 2019; Schmiedel and Lehner, 2019) and `D+M` (Section S2.3.2).

Specific epistasis among *enriched* protein contacts—position pairs which are both significantly enriched after correcting for multiple testing (Benjamini and Hochberg 1995, *α*_*BH*_ = .1) and within 5Å in the crystal structure—is disproportionately positive, as first reported in Diss and Lehner (2018) (Fig. 4c). In Fig. 4, we highlight an extreme example—positions L4 and L4 in Fos and Jun, respectively—for which all of the ≈ 30 amino acid interactions with *p*-values below *α* = .05 are positive in each replicate (Fig. S4a). The predominance of positive interactions is likely, in part, explained by two considerations: the majority of mutations at positions engaged in protein-protein contacts are deleterious in isolation (Fig. 5a), and R&R is relatively underpowered to detect negative interactions among the most deleterious mutants (Section 3.1). The latter limitation is not unique to R&R, as a non-rank based method, `D+M`, produces a similar bias and is also consistent with pervasive positive epistasis among L4/L4 mutations (Fig. S9). Detection of overwhelming positive epistasis among mutations at the L4/L4 pair persists when a two-sided *p*-value is computed, suggesting that methodological biases cannot fully explain this result (Section S2.3).

Positions T8 Fos and V8 Jun present a notable exception to the widespread positive epistasis among protein contacts (Fig. 4a-e). The vast majority (≈95%) of significant interactions between T8 and V8 are negative in each replicate (Fig. S4b), a result that cannot readily be explained by methodological biases nor technical artifacts (see Section S2.3). The majority of single mutations of T8 Fos are neutral or beneficial in isolation, with on average 14 out of 20 mutants exhibiting fitness greater than the wildtype sequence across replicates, while the V8 Jun mutants span a larger fraction of the measurement range.

#### PDZ3-CRIPT

Sequencing coverage in the PDZ3-CRIPT dataset was lower than for that of Fos-Jun, resulting in less precise fitness measurements. Noisier fitness measurements likely contribute to a lower Spearman’s correlation between single mutant rank and background-averaged rank in PDZ3-CRIPT compared to Fos-Jun (Fig. 5d, 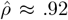 versus .99 in each replicate, respectively, and see Section S2.5.3). In particular, overestimation of single mutant fitness values due to low initial read counts may have resulted in spurious detection of negative SE (Section S2.5.3).

To be conservative in the detection of enriched position pairs, we chose the significance threshold in each block that maximized contact accuracy and recall (Fig. S14b, *α*_1_ = .034, *α*_2_ = .021). As in Zarin and Lehner (2024), we also identify position pairs enriched for *positive* interactions using the same *p*-value thresholds, respectively. While the pairs most enriched for positive interactions are within 5Å—partly by design—several position pairs in block 1 overtly contravene this trend. Specifically, S-1 CRIPT and L342 and G345 in PDZ3, and T-2 CRIPT and G330 PDZ3 at distances of approximately 8, 11, and 10Å, respectively, are significant after correction for multiple testing (*α*_*BH*_ = 10^−2^; Fig. 4g,h). These position pairs were previously identified as “specificity-changing mutations” in Faure and Lehner (2024) and/or implicated in allosteric regulation in PDZ3 binding (Raman et al., 2016). In isolation, both T-2 CRIPT and G330 PDZ3 harbor many of the most deleterious mutations (Fig. 4i). However, a subset of amino acid pairs at these positions results in appreciable fitness gains (Fig. 4i), though with absolute fitness still below the wildtype sequence (Fig. S15). In contrast, S-1 CRIPT and L342 and G345 PDZ3 single mutants have more modest effects on fitness, and while detected SE is on average positive, several amino acid pairs exhibit negative SE (Fig. 4i and see Fig. S15).

H372 PDZ3 and T-2 CRIPT present the largest number of “strong” interactions among amino acid pairs (Fig. 4i and Fig. S15). Single mutations at both of these positions are very deleterious, with almost all variants in the bottom quartile. In some cases, the test statistic approaches or reaches its maximum value (Fig. 4i). Though, as with the G330 PDZ3/T-2 CRIPT pairs, epistasis does not fully recover wildtype fitness (Fig. S15). As the ranks of single mutations at H372 and T-2 are similarly deleterious, and many amino acid pairs exhibit large, positive rank deviations, R&R may underestimate the proportion of non-zero interactions (Fig. 4i). For example, the *p*-values associated with H72V and T-2R are above *α* = 0.05 in each replicate. However, the observed rank deviations are unlikely to be explained by experimental noise, as suggested by the corresponding meta *p*-value 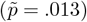.

### 3.3 Model misspecification leads to spurious detection of specific epistasis

Finally, we analyze a comprehensive DMS of 55 of the 56 amino acids in the immunoglobin fragment G (IgG) binding domain of protein G, GB1 (Olson et al., 2014). Our primary interest in the DMS of GB1 is that single-trait GE is *a priori* thought to be insufficient to explain GB1-IgG binding. Instead, prior work supports a three-state equilibrium model, where the protein’s folding *and* binding energies govern transitions among unfolded, folded, and ligand-bound states (Otwinowski, 2018; Nisthal et al., 2019). In this case, fitness—a proxy for binding affinity—is a bivariate nonlinear function of two latent additive traits, Λ^fold^ and Λ^bind^ (Eq. S8). The presence of two latent traits implies that the results of Section 2.1 no longer universally hold, and the ranks of mutations will not necessarily be preserved across backgrounds. Indeed, the contact accuracy of R&R at the level of position-pairs is substantially lower than for Fos-Jun (Fig. S12b), though R&R performs comparably to or slightly below prior, more heuristic procedures (Section S2.4). When `D+M` is used to fit a three-state model, contact prediction accuracy improves appreciably (Fig. S12). As such, the DMS of GB1 presents an opportunity to evaluate the consequences of model misspecification, in the form of an additional latent trait, on R&Rs detection of SE in an empirical data set.

A key feature of the three-state model is that a mutation *i* with an adverse effect on binding is more deleterious in the background of an unstable mutant *j*. In this scenario, the rank of mutant *i* in the background of *j* will likely be lower than expected given its single mutant rank, i.e., 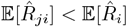, resulting in a preponderance of negative 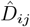 values and spurious negative SE among all such pairs. The converse holds when a stabilizing mutation *i* occurs in the background of a mutant *j* with adverse effects on folding: 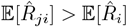 yielding 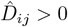 and spurious positive SE.

We therefore suspected that a mutation’s *residual*, defined as the difference between its rank based on all possible backgrounds and single mutant rank (Fig. 6c), would correlate with its effects on folding energy, 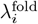. Indeed, variants with adverse effects on 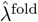—independently measured by Nisthal et al. (2019)—exhibit more negative residuals relative to more stable variants (Fig. 6b and Fig. 6d, linear fit, *R*^2^ = .26), larger numbers of significant associations, and higher pseudo-false positive rates (Fig. 6f, lower triangle). Indeed, 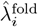 is a much better predictor of pseudo-false positive rate than single mutant phenotype (Fig. 6f, upper triangle). To understand this phenomenon, we reproduce Fig. 1b of Otwinowski (2018), in which the inferred folding and binding energies of each GB1 mutant are plotted with respect to the wildtype sequence (Fig. 6e). Here, we observe that variants with estimated adverse effects on folding (and thus 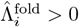) *and* fitness approximating the wildtype fitness exhibit the highest pseudo-false positive rates; in Fig. 6e, the size of each point is proportional to the number of false positives. Simulations under the assumption of a three-state model without SE qualitatively reproduce these results (Section S2.4.1).

**Figure 6.**
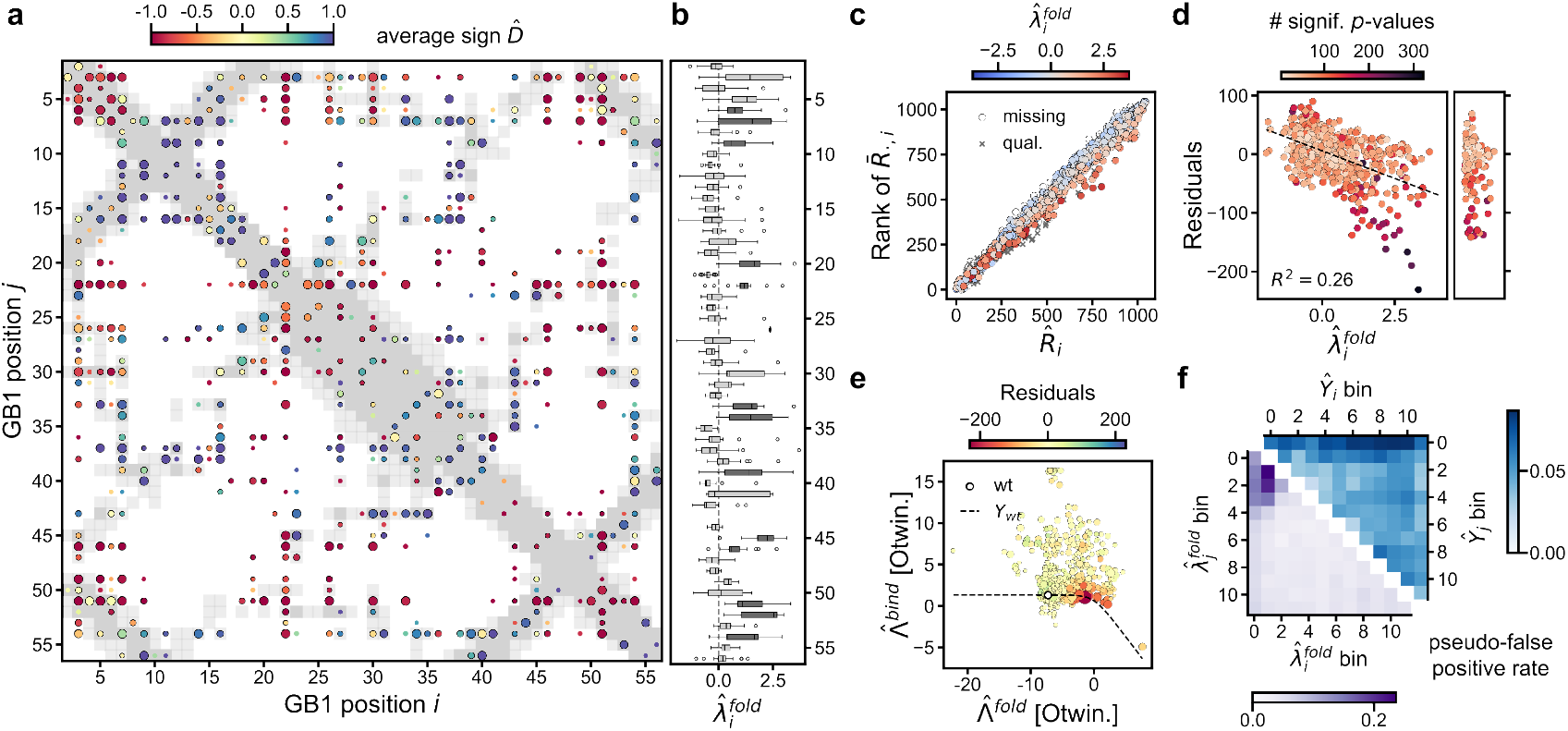
Model misspecification results in spurious detection of epistatic effects. **(a)** All position pairs enriched for significant epistatic interactions (*α* = 10^−1^) after multiple testing correction (Benjamini and Hochberg 1995, *α*_*BH*_ = 10^−1^) in GB1 are shown as colored points, with color indicating the average sign of the significant interactions. The size of the point is proportional to the −*log*_10_ enrichment *p*-value, with pairs significant at a false-discovery rate (FDR) of *α* = 10^−2^ outlined in black. All position pairs within 5 and 8Å are denoted by dark and light gray filled boxes, respectively. **(b)** Box plots of the folding energies of GB1 single mutants, 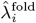 estimated independently in Nisthal et al. (2019), for each position. Positions with significantly greater average folding energies (one-sided T-test, FDR *α* = 10^−1^) are colored in dark gray. **(c)** The rank of the average rank of each mutation across all backgrounds, 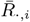, plotted as a function of single mutant rank 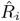. Each point is colored by 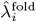, except for mutants with missing measurements, denoted in light gray, and mutants with “qualitative” measurements in dark gray (see Nisthal et al. 2019). **(d)** Residuals, defined as the difference between the *y* and *x*-axis values of **c**, as a function of 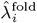. The dotted line denotes a linear fit, estimated from all of the quantitative measurements. The residuals for the qualitative measurements are shown to the right. Each single mutant is colored by the number of significant *p*-values across all double mutants containing that mutation (*α* = 10^−1^). **(e)** Figure 1b of Otwinowski (2018) reproduced, with each single mutant additionally colored by its residual (*y*-axis in **d**). The size of the point corresponds to the number of pseudo-false positives involving each mutant, where pseudo-false positives are defined as significant associations (*α* = 10^−1^) between positions at distances greater than 8Å. **(f)** The pseudo-false positive rate (*fpr*) is shown as a function of the binned ranks of the single mutants (top right) and 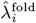 (Nisthal et al. 2019, bottom left).

## 4 Discussion

Quantifying epistasis usually requires specification of the measurement scale. Fisher (1919) introduced the concept of “epistacy” in the context of quantitative traits as a deviation from additivity. In evolutionary biology, SE is usually defined as a deviation from a multiplicative fitness map (Phillips, 2008; Araya et al., 2012). More recently, researchers have defined SE as a deviation from a model of GE fitted under minimal assumptions about the form of the nonlinearity, *g*. For example, modeling *g* as a sum of monotonic functions, such as spline functions or sigmoids (Otwinowski et al., 2018; Faure and Lehner, 2024). Non-parametric procedures, such as Zhou and McCandlish (2020), Zhou et al. (2022), and Aghazadeh et al. (2021), impose even fewer constraints on scale, but in not explicitly accounting for global nonlinearities may over-estimate the prevalence of SE.

By redefining epistasis in the context of rank-statistics, R&R entirely circumvents the choice of scale. Our work follows from the observation that, as long as *g* is monotonic—but not necessarily nonlinear—SE can be detected as a deviation from a given reference order. In a combinatorial DMS, the fitness values of the single mutants specify a valid reference order. In the absence of measurement noise, and when *g* is strictly increasing (or decreasing), the relationship between a given rank deviation and the direction and magnitude of an interaction *λ*_*ij*_ is immediate (let *σ*_*ϵ*_ = 0 in Eq. 4). Measurement noise, of course, obscures SE.

We contribute a principled hypothesis testing procedure, R&R, that accounts for heteroskedasticity in the test statistic due to variation in (1) the slope of the nonlinearity, (2) the density of single effects on the latent trait Λ, and (3) variance of the fitness estimates conditional on Λ. To “learn” the distribution of 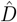 as a function of the ranks of the constituent mutations, R&R generates an ensemble of synthetic datasets under the assumption of an error model—the only context-specific and parametric element of R&R. In the analyses of sequencing-based protein DMSs presented here, we employed a Poisson error model. This choice allowed R&R to operate on minimally processed variant count data. However, additional post-processing and alternative error models, e.g., normally distributed errors, would allow R&R to combine information across replicates and account for batch effects, as observed in the PDZ3-CRIPT dataset and accounted for in Zarin and Lehner (2024). In the future, R&R could be combined with more sophisticated error models and simulation frameworks, for example, that of Rao et al. (2024). Superior modeling of the error distribution would likely improve R&Rs statistical power; excess variance in the bootstrap sample hindered R&Rs ability to detect weak SE in simulations.

Despite its simplicity, R&R accurately identifies true SE and protein contacts in simulated and empirical data sets, respectively, performing comparably to (or slightly better than) a neural network based procedure, `D+M`, on the task of contact prediction in a DMS of Fos-Jun. The application of R&R to a diverse set of DMSs—with substantial variation in measurement precision—demonstrated its robustness, while also revealing several limitations:

The test statistics for a given mutant *i* are correlated across variants *j* as 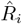 is a constituent of 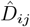 for all *j* ≠ *i*. If the estimated single mutant rank is much higher than the true rank, 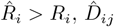 will tend to underestimate its true value on average (and vice versa for an underestimated single mutant rank). The same problem arises in standard estimation of epistatic coefficients, but is mitigated by estimation via linear regression where fitness estimates for higher order mutants contribute to estimation of lower order effects (Dupic et al., 2024). When the fitness values of single mutants are estimated with high precision—as is the case in numerous DMSs—errors in the single mutant ranks will make minimal contributions to the false positive rate. However, when single mutant ranks are estimated with non-negligible error—as in the PDZ3-CRIPT data set—we expect to, and likely do, detect spurious SE. Alternative definitions of the test statistic could potentially mitigate susceptibility to errors in the single fitness estimate. For example, average rank with respect to a subset of mutant backgrounds could determine the reference order.

R&R implicitly assumes that the estimated double (and single) mutant fitness values are independent and identically distributed conditional on their latent trait values. In practice, mutants are present at varying frequencies in the pre-selection pool, resulting in variation in measurement precision. Imprecise fitness measurements will increase the volatility of the bootstrap sample, reducing power to detect SE among more precise fitness measurements. To mitigate this bias, one could potentially integrate over uncertainty in the double mutant rank when computing the test statistic or devise a more sophisticated bootstrapping scheme that accounts for variation in measurement precision.

R&R relies weakly on the assumption that the second order terms, *λ*_*ij*_, are sparse. Under this assumption, the empirical distributions will approximate the null distributions of the test statistics. When SE is not sparse, R&R will detect the largest interactions, underestimating the complexity of the genotype-to-phenotype map. However, even when SE is sparse, effects may not be uniformly distributed across the measurement range. Such a scenario may arise due to the fact that mutations at the same position tend to have similar fitness values, *Y*_*i*_, and will also tend to interact with similar partners—as observed with H372 in PDZ3 and T-2 in CRIPT. This will reduce power to detect weaker positive interactions for positions in the single rank neighborhoods of H372 and T-2 mutants. More generally, the power to detect significant interactions depends on the density of SE in a mutant’s single effect neighborhood.

We focus on DMSs where the fitness values of both single and the majority of double mutants have been observed. The large number of double mutants allows R&R to estimate the empirical distributions of the rank-indexed test statistics. Many DMSs, however, more sparsely sample the sequence space and include higher-order mutants. For example, “pathway” DMSs assay a combinatorially complete set of mutants that interpolate between two sequence endpoints (e.g., Weinreich et al. 2006; Wu et al. 2016; Poelwijk et al. 2019; Moulana et al. 2023). Extending R&R, and applying rank-based procedures for detecting interactions more generally (see Crona et al. 2017; Lienkaemper et al. 2018) to these more common, sparser datasets may provide additional insights into epistasis across proteins.

Beyond details of the system under study—namely, the distribution of single fitness effects, the shape of the nonlinearity, and the magnitude of measurement noise—the proposed rank-based procedure imposes inherent constraints on power to detect SE in different parts of the measurement domain. For example, it would be impossible to detect strong negative SE between deleterious mutations, or to likewise detect strong positive SE among the most beneficial mutations. More generally, the magnitude of 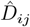 is constrained by the single mutant ranks of the constituent mutations. This being said, fitness estimates at the bounds of the measurement range are perhaps more likely to be subject to technical artifacts, such as non-specific binding at the low end (Olson et al., 2014), or PCR amplication biases at the higher end (Faure et al., 2020). In this case, lack of power to detect SE among mutants at the respective extrema of the measurement range is a feature rather than a limitation.

More profoundly, analyzing DMSs from a rank-based perspective exposes systematic biases in the detection of SE—with R&R or other procedures—across the measurement range. For example, both R&R and `D+M` are underpowered to detect SE, and particularly negative SE, among deleterious mutations. Our ability to derive general principles about proteins from distinct DMSs requires an understanding of how experimental and statistical biases influence the distribution of SE detected in a given experiment. For example, to conclude that SE is on average positive or negative, necessitates a proper accounting of statistical power to detect SE of one sign versus the other.

In the present work, we have arguably treated the nonlinearity *g* as a “nuisance”, which conceals the “true” epistatic landscape. When *g* is induced entirely by experimental design, for example, by a lower detection threshold, this treatment is uncontroversial. However, when *g* emerges from the physics of the protein, one can not necessarily neatly distinguish between a global nonlinearity and direct physical interactions (see Dutta et al. 2018). For one, recent work demonstrates that GE can emerge from numerous microscopic interactions among positions (Lyons et al., 2020; Reddy and Desai, 2021). This finding reveals a fundamental ambiguity in the specification of genotype-to-phenotype models: Is a GE model—where the latent additive trait includes sparse higher-order terms—to be privileged over a dense model, with epistatic interactions at many orders, that does not explicitly account for global nonlinearities (e.g., see Dupic et al. 2024; Zhou et al. 2022)?

Our work presents immediate areas for future research. While monotonicity is common in biology, it is by no means a rule. In addition, as we demonstrated in Section 3.3, the presence of an additional latent trait(s) results in spurious inference of SE. Extending rank-based detection of SE (and assessment of GE) to higher dimensional latent spaces represents a rich area for future work.

## 5 Materials and Methods

### 5.1 Datasets

#### 5.1.1 G protein binding domain

In Olson et al. (2014), the authors generated an almost complete library of the single and double mutants of 55 of 56 residues in protein G domain B1 (GB1). The binding affinities of all mutants to IgF fragment were measured in a sequencing-based fitness assay. As the selection assay was repeated three times using the same input library, we summed the reads from all three replicates to estimate fitness. After removing variants with fewer than 21 reads in the initial pool, all single mutants (*n* = 55 *×* 19 = 1,045) and 97% of double mutants (*n* = 519,030) remained.

#### 5.1.2 Fos-Jun heterodimer

In Diss and Lehner (2018), the authors assayed the binding affinity of (1) all single mutants in the Fos and Jun proteins; and (2) the majority of possible *trans* double mutants between the Fos and Jun proteins. The phenotypic assays were performed in triplicate, with three separate input and post-selection libraries.

Raw sequencing reads were downloaded from the Sequence Read Archive (SRR5952429-SRR5952434). The sequencing data were first filtered to remove reads for which (1) the adapter sequences were more than two bases removed from the designed adapter; (2) the unique molecular identifier had appeared earlier in the same sample; or, (3) the read contained an “N”. The filtered reads were translated from nucleotide to amino acid sequences and aligned to the wildtype sequences of Fos and Jun to generate read counts for each single and double mutant.

In each replicate, variants with fewer than 11 reads in the input pool were set to missing. In contrast to Diss and Lehner (2018), variants were not filtered on read counts in the post-selection pool, resulting in the inclusion of, on average, 43% 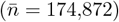 of all double mutants across the three replicates and all single mutants (*n* = 1, 280).

#### 5.1.3 PDZ3 and CRIPT binding

We analyzed a subset of the data collected in Zarin and Lehner (2024) consisting of all single and a fraction of the double *trans*-mutants of the 100 amino acid region (amino acids 303-402) of the third PDZ (**P**ostsynaptic density 95; **D**rosophila disc large tumor suppressor; **Z**onula occludens-1) domain of PSD-95 and its cognate ligand CRIPT (*L* = 8 amino acids). Mutants in the first and latter 50 amino acids of the PDZ3 region were assayed in separate batches, resulting in pronounced batch effects (see Zarin and Lehner 2024 and Section S2.5). Therefore, we analyzed the two halves of the 100 amino acid region, referred to as blocks 1 and 2, separately. In addition, we removed all positions for which the average single mutant read coverage was below 20. In each block, 43 positions remained (positions 303-345 and 353-395).

In each of the three replicates, variants with fewer than 11 input reads were set to missing. Subsequently, variants with greater than 95% missing data across all double mutants were removed. After filtering, fitness values were reported for 96% of CRIPT mutants (*n* = 154), 92% (*n* = 791) of single PDZ3 mutations and 66% (*n* = 91, 017) of *trans*-double mutants in block 1, and 98% (*n* = 840) single PDZ3 mutations and 89% (*n* = 122, 887) of *trans*-double mutants in block 2, on average, respectively, across the three replicates.

### 5.2 Testing for specific epistasis

#### 5.2.1 Computing the test statistic

Double mutants necessarily involve two mutations at distinct positions. If there are *L* positions in the protein and *M* possible states, one can only ever hope to observe *L*_*d*_ := (*L* − 1) *×* (*M* − 1) double mutants in a given background. Thus, to compute the test statistic 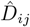 (Eq. 6), we adjust the single rank of *i* by excluding all mutations at position *j* (and vice versa), where we have abused notation in using *i* and *j* to refer to both position and amino acid mutant.

As fewer mutants may be observed in some backgrounds due to insufficient coverage in the initial sequencing pool, all of the ranks are transformed onto the same scale such that the ranks in a each background range from 0 to *L*_*d*_ − 1 (Section S1.2.1). Tied values are resolved using mid-ranks (Section S1.1).

In the PDZ3-CRIPT data set, where the number of PDZ3 mutations far outnumbered that of CRIPT, we employ a reweighted test statistic (Section S1.3).

#### 5.2.2 Poisson error model

The pre- and post-selection read counts in the *b*-th bootstrap replicate 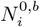 and 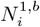 for mutant *i*, respectively, are assumed to be Poisson distributed with means specified by their observed value, 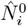 and 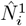,

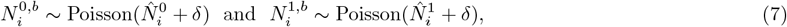

where *δ* = 1 is a pseudocount. The fitness estimate for mutant *i* in the *b*-th replicate is then given by,

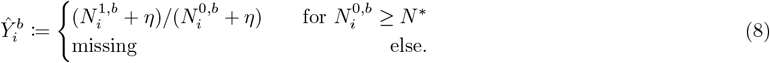

where 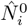 is the observed initial read count and *N*^*^ is a minimum initial read count threshold, and likewise for double mutants. While *δ* > 0 biases the bootstrap sample, inclusion of a pseudo-count in Eq. 7 introduces randomness into the fitness measurements of variants with zero post-selection counts.

#### 5.2.3 Estimating the *p*-value of mutant interactions

The bootstrapping procedure described in Section 2.2.2 is used to generate an ensemble of test statistics for each pair of double mutant ranks, *r* and *s*. The empirical distribution of 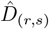 is then given by,

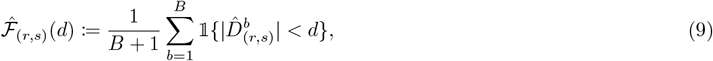

where 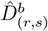 is computed from Eq. 6 for the mutants ranked *r*^th^ and *s*^th^ in the *b*^th^ simulation. As certain combinations of mutant ranks are not observed in a given bootstrap replicate, the matrix 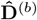 is imputed using a nearest neighbor approach (Section S1.2.2). The *p*-value for a pair of mutations *i* and *j* with estimated ranks *r* and *s*, is then given by,

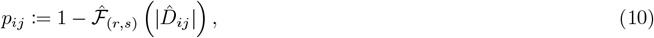

where 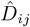 is the observed test statistic.

#### 5.2.4 Combining results across replicates

When experimental replicates are available, as in the Fos-Jun and PDZ3-CRIPT data sets, we use Fisher’s method (Fisher, 1925) to estimate a single *p*-value for each position-amino acid pair, 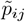. In the event of *R* replicates,

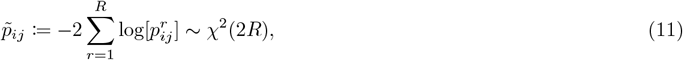

and 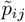 is *χ*^2^ distributed with 2*R* degrees of freedom under the assumption of uniform *p*-values in each replicate *r*. To reduce false positives, we exclude all mutant pairs that were observed in only a single replicate, i.e., *R* ∈ {2, 3}.

#### 5.2.5 Testing for specifically epistatic position-pairs

A natural estimator for the strength of the interaction between two positions *i* and *j* is to combine the amino acid level *p*-values. Letting *k* and 𝓁 refer to the amino acid mutation at positions *i* and *j*, respectively,

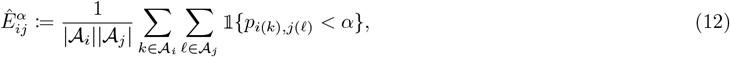

where 𝒜_·_ and |𝒜_·_| are the set and size of the set at a given position, and *α* is a fixed *p*-value threshold. Following previous work, 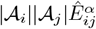 was assumed to follow a hypergeometric distribution, parameterized by the total number of position pairs and the total number of *p*-values less than *α* across all such pairs (e.g., Schmiedel and Lehner 2019; Faure and Lehner 2024). All mutant pairs involving null alleles, i.e., premature stop codons, were excluded from the enrichment analysis as these mutations should *a priori* not engage in SE. Where replicates were available, we use the meta *p*-values to conduct the enrichment test (Eq. 11).

In the main text, *α* = .05 in the Fos-Jun analyses, *α* = .034, .021 in blocks 1 and 2 of PDZ3-CRIPT analyses, and *α* = .1 in the GB1 analysis, respectively. In Sections S2.3.2, S2.4 and S2.5.1, we conduct the enrichment tests over a range of *α* values.

#### 5.2.6 Method comparison

`DiMSum` (Faure et al., 2020) was applied to simulated and empirical data sets after filtering on initial read counts, to estimate the mean (log) relative fitness values and their standard errors under a Poisson error model. The estimates from `DiMSum` were used as input to `MoCHI` (Faure and Lehner, 2024) to fit a GE model, where the nonlinearity was assumed to take the form of (1) a sum of arbitrary sigmoid functions (GB1-like simulations and Fos-Jun), (2) a two-state thermodynamic model (GB1), and (3) a three-state thermodynamic model (GB1). See Section S1.4 for more details.

### 5.3 Simulations

#### 5.3.1 Single-trait global epistasis

We simulate a DMS under the assumption of a two-state thermodynamic model, as described in Otwinowski (2018). The underlying additive trait, Λ, corresponds to the binding energy of the protein, and the Boltzmann distribution determines the proportion of bound protein,

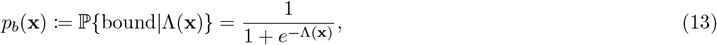

where Λ(**x**) is defined as in Eq. 1.

In a DMS, protein function is often not measured directly. Rather, a library of protein variants is first constructed, often in bacteria or yeast. The variant library is sequenced in the presence and absence of a selection pressure tied to protein function. The enrichment of variants, as measured by sequencing, is then a proxy for the protein function under study. The estimated fitness *Ŷ*_*i*_ of a variant *i* is given by,

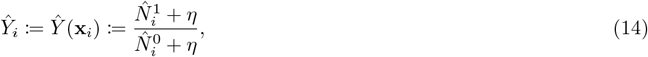

where 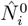 and 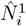 are the number of reads attributed to mutant *i* before and after selection, and *η* = 1.

To sample the post-selection reads, 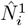 and 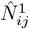, we adopt the likelihood model of Otwinowski (2018), which permits a non-zero probability of non-specific binding, *δ*. In the selective binding assay of Olson et al. (2014), the number of molecules after selection, 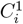, can be approximated as,

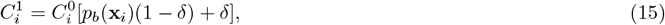

where *p*_*b*_(***x***_*i*_) is defined in Eq. 13, and 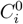 is the number of mutant *i* cells in the initial pool. We assume that both 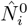 and 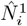 are Poisson random variables, independent conditional on 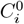,

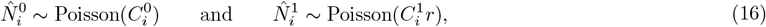

where *r* is a global factor that accounts for differences in the number of cells and sequencing reads before and after selection.

#### 5.3.2 Specific epistasis

Motivated by empirical results, we assume that the strength of interactions between amino acids *k* and 𝓁 at two positions *i* and *j, λ*_*i*(*k*),*j*(𝓁)_, is inversely related to their physical distance in the crystal structure *d*_*ij*_, where *d*_*ij*_ is measured as the minimum Euclidean distance between the heavy atoms comprising the native amino acids at both positions. Specifically, when *d*_*ij*_ ≤ 5Å, *λ*_*i*(*k*),*j*(𝓁)_ are *iid* random variables for all amino acid pairs *k* and 𝓁, and when *d*_*ij*_ > 5Å, *λ*_*i*(*k*),*j*(𝓁)_ is set to zero:

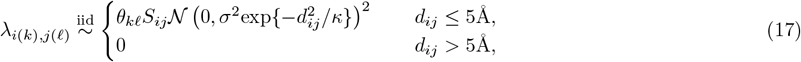

where, *S*_*ij*_ ∈ {−1, 1} is an unbiased signed Bernoulli, and *κ* and *σ* are two scaling parameters which control the decay of effects with physical distance and the effect magnitude, respectively. In addition, we introduce a Bernoulli random variable *θ*_*k*𝓁_ to sparsify the non-zero coefficients, reflecting the fact that often only a subset of amino acid combinations among epistatic position pairs exhibit non-zero interactions. We assume that 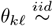 Bernoulli(*θ*), for *θ* = .5. In addition, all values of *λ*_*i*(*k*),*j*(𝓁)_ *<* 10^−1^ are set to zero.

#### 5.3.3 Simulation scenarios

Under the assumption of a two-state thermodynamic model, we consider the following simulation scenarios:

1. **Equal initial frequencies**. The initial cell counts (scaled by the initial read counts) are shared among single and double mutants, with 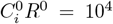 and 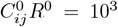, respectively. Initial read counts are sampled from Poisson distributions parameterized by the scaled cell counts (Eq. 16). The latent trait value for the wildtype sequence is set to its inferred value from Otwinowski (2018), Λ_*wt*_ = 1.34, and single mutant effects on Λ are sampled from a uniform distribution, i.e., *λ*_*i*_ ∼ Unif[−3.5, 3.5] for all mutants *i*. The Poisson model (Eq. 16) is further parameterized by *η* = 1, *δ* = 0, and *r/R*^0^ = 4. Epistatic coefficients *λ*_*ij*_ are sampled from Eq. 17 with *σ*^2^ = 2, *κ* = 2, and *θ* = .5. We run an additional simulation, *1b*, in which we reduce the read counts by a factor of ten.
2. **Cell frequency variation**. To introduce additional variation in the sequencing read counts, we assume that 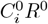 and 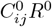 are sampled with replacement from the empirical distribution of read counts in the GB1 data set (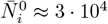 and 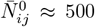). To limit the number of samples with initial read counts below a threshold *N*^*^ = 21, we set all empirical read counts to *N*^*^ prior to sampling. All samples with sampled read counts below *N*^*^ are then dropped from the analysis. All other procedures and parameters are inherited from *Simulation 1*.
3. **Non-specific binding**. All parameters are inherited from *Simulation 1* except a small probability of non-specific binding *δ* (Eq. 16) was introduced. We let *δ* = 7 · 10^−4^, as estimated in Otwinowski (2018).
4. **Realistic** Λ **distribution**. We sample the read counts as in *Simulation 2*. However, we now let the energy estimates of Otwinowski (2018) specify the single mutant effects on Λ. To reduce the number variants with post-selection reads 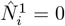, and as parameter estimates for very deleterious mutants are likely to be inaccurate, we resampled the effects for mutants with *λ*_*i*_ ≥ 4 uniformly on the interval [0, 4]. As in *Simulation 3*, we allow for non-specific binding, with *δ* = 7 · 10^−4^.

We run an additional simulation, *4b*, in which we reduce the single and double mutation read count distributions by a factors of 10 and 5, respectively, and reduce the minimum read count threshold to *N*^*^ = 11 To reduce sample dropout we set all empirical read counts below *N*^*^ to *N*^*^.

Finally, we simulate data under the assumption of a three-state thermodynamic model, parameterized by the analysis pf the GB1 data (Olson et al., 2014) by Otwinowski (2018). In this model, two latent traits, corresponding to a variant’s folding and binding energies, determine fitness (see Section S2.4.1 for details).

## 6 Data and software availability

All variant count files and code required to reproduce the analyses are available at github.com/marync/resample and reorder. The raw sequencing reads associated with each empirical data set were previously made available to the public.

## 7 Acknowledgments

We thank Federica Ferretti, Alisdair Hastewell, Nikos Ignatiadius, Rama Ranganathan, Abigail Skwara, Rebecca Willett, and attendees of the NITMB research-in-progress for helpful conversations. We thank Taraneh Zarin for providing guidance with respect to the PDZ3-CRIPT data. MC was supported by NIGMS of the NIH under award number R35GM151211 and an NITMB fellowship supported by grants from the NSF (DMS-2235451) and Simons Foundation (MP-TMPS-00005320). All simulations and analyses were performed on the Midway cluster, supported by the Research Computing Center at the University of Chicago.

## 8 Author contributions

MOC, BLA, YBS conceptualized and designed the research; MOC performed the analyses; MOC, BLA created the figures; MOC wrote the paper; MOC, BLA, YBS revised the paper.

