## Supporting Information for "Robust detection of specific epistasis using rank statistics"

### Supporting information for
#### Robust detection of specific epistasis using rank statistics

Maryn O. Carlson[1*], Bryan L. Andrews[2,3,4*], and Yuval B. Simons[5*]

[1]National Institute for Theory and Mathematics in Biology, Chicago, IL, USA
[2]Department of Biochemistry and Molecular Biology, University of Chicago, Chicago, IL, USA
[3]Center for the Physics of Evolving Systems, University of Chicago, Chicago, IL, USA
[4]Center for Living Systems, University of Chicago, Chicago, IL, USA
[5]Section of Genetic Medicine, University of Chicago, Chicago, IL, USA

April 8, 2025

### S1 Supplementary methods

We provide additional methodological details for Resample and Reorder (R&R), and its application to data.

#### S1.1 Accounting for ties

Fitness measurements from sequencing-based assays are often discrete. In the experiments analyzed herein, a fitness measurement is given by the ratio of post- and pre-selection read counts, $N^1$ and $N^0$, respectively,

$$\hat{Y} \coloneqq (N^1 + \eta)/(N^0 + \eta) \in \mathbb{N}_0 \times \mathbb{N}_0,$$

where, $\eta = 1$ is a pseudo-count. As a consequence, there is a non-negligible probability that distinct variants with similar true fitness values may have the same estimated fitness values, particularly when $N_0$ is small. Tied fitness measurements are particularly common for low fitness variants with true fitness values near the limit of detection, as $N_1$ may be equal to zero for these variants. (The limit of detection is determined by features of the experiment, such as sequencing depth and ligand concentration.)

We use mid-ranks (Kendall, 1945) to account for ties in the estimated fitness values prior to estimating the test statistic $\hat{D}$. That is, we assign to the tied measurements the average of the corresponding ranks. For example, if $n$ mutants have estimated fitness values equal to the minimum fitness value, all $n$ mutants are assigned a rank of $(n-1)/2$ (assuming zero indexing).

In the bootstrap sample, however, we resolve ties among single mutant fitness estimates by random sampling in order to reduce the correlation between bootstrap replicates and to associate each mutant with a unique rank. (We use mid-ranks to resolve ties among double mutants in the bootstrap samples as described above.)

#### S1.2 Accounting for missing data

##### S1.2.1 Transforming ranks on the same scale

Consider an $L$ length protein with $K$ amino acid states. (In most cases, $K = 20$.) The single mutant ranks are defined on the set $\{0, \dots, M-1\}$, where $M = L \times (K-1)$. In a given background, $i$, the phenotypes of at most $M - (K-1)$ mutants are observed as it is impossible to make two mutations at a single site. In addition, some double mutants may not be observed due to insufficient coverage in the initial sequencing pool. Let $M_i < M$ be the number of double mutants observed in background $i$. To map ranks on the set $\{0, \dots, M_i - 1\}$ to $\{0, \dots, M-1\}$, we apply a linear transformation,

$$\tilde{R}_{im} \to \tilde{R}_{im}\left(\frac{M-1}{M_i-1}\right) =: \hat{R}_{im}, \tag{S1}$$

where $\hat{R}_{im}$ denotes the "interpolated" rank. Interpolated ranks are used in the computation of $\hat{D}_{im}$, but the double mutant ranks are scaled to the maximum number of observed double mutants.

##### S1.2.2 Imputing the bootstrapped test statistic matrix

As noted above, it is impossible to observe two different mutations at the same position. Thus, even in a complete data set, the test statistic in a given bootstrap sample $b$, $\hat{D}^b_{(r,s)}$, may not be estimated for some set of $r$ and $s$ rank pairs observed in the data. To ensure that the distributions of $\hat{D}_{(r,s)}$, $\hat{\mathcal{F}}_{(r,s)}$, are estimated from approximately the same number of simulations, we impute the matrix of $\hat{D}^b_{(r,s)}$ values using an iterative $k$-nearest neighbors algorithm within each bootstrap replicate $b$:

For a given missing data point $(r', s')$, we sample one value from among its $k = 1$ nearest neighbors, defined by the set of indices $\{r'-1, r'+1\} \times \{s'-1, s'+1\}$. If all values in the 1-neighborhood of the missing value are also not observed, a value is sampled from the next $k$-neighborhood, defined by the set of indices $\{r'-k, \dots, r'-1, r'+1, \dots r'+k\} \times \{s'-k, \dots, s'-1, s'+1, \dots s'+k\}$. This procedure is repeated until a neighborhood with a non-missing value is found, or until $k = 5$.

#### S1.3 Computing the test statistic

In practice, when computing the test statistic, $\hat{D}_{ij}$, we modify the single mutant ranks to exclude all mutations at a given position $i$. (In an $L$-length protein with $K$ states, we will observe at most $L - 1 \times K - 1$ mutations in a given background.) We refer to this adjusted rank of mutation $j$ in the background of $i$ as $\hat{R}^{-i}_j$ and define the test-statistic more precisely as,

$$\hat{D}_{ij} = \hat{R}_{ij} - \hat{R}^{-i}_j + \hat{R}_{ji} - \hat{R}^{-j}_i, \tag{S2}$$

where we have abused notation in using $i$ and $j$ to refer to both mutations and positions.

The range of possible rank deviations, i.e., differences in the double and single mutant ranks, $\hat{R}_{ij} - \hat{R}_j^{-i}$, is determined by the number of double mutants observed in a given background. In the PDZ3-CRIPT study (Zarin and Lehner, 2024), differences in protein and ligand length ($L = 43$ and $L' = 8$, respectively) imply that the test statistic will be dominated by the rank deviations of the PDZ3 mutants. We therefore propose a weighted test statistic, $\tilde{D}_{ij}$,

$$\tilde{D}_{ij} = \alpha(\hat{R}_{ij} - \hat{R}_j^{-i}) + \beta(\hat{R}_{ji} - \hat{R}_i^{-j}), \tag{S3}$$

where $\alpha, \beta > 0$, where $\hat{R}_j^{-i}$ and $\hat{R}_i^{-j}$ were defined above. For example, when $\alpha = L'/L$ and $\beta = 1$, the rank deviations are transformed onto the same scale. Eq. 6 is given by $\alpha = \beta = 1$.

#### S1.4 Method comparison

In the main text, we compare R&R to a procedure for detecting SE outlined in Zarin and Lehner (2024). The procedure consists of two steps. First, `DiMSum` (Faure et al., 2020) is used to estimate the mean log (relative) fitness value, $\hat{Y}^{\mathtt{D}}$, and its standard errors, $\hat{s}$, of each variant from its pre- and post-selection read counts. The standard errors are estimated under the assumption of a Poisson read model, with additional replicate specific terms to account for over-dispersion and batch effects. In the case of a single replicate, the error model reduces to $\hat{s}_{ij}^2 := 1/N_{ij}^0 + 1/N_{ij}^1$, and corresponds (approximately) to the "Poisson error model" described in Section 5.2.2. Second, `MoCHI` (Faure and Lehner, 2024), is used to fit a GE model of a pre-specified class (e.g., two-state thermodynamic model, sum of sigmoid functions) to the estimated fitness values (and their standard errors). Unlike `DiMSum`, `MoCHI` assumes a normal error model, with the mean and variance of each fitness estimate specified by their `DiMSum` estimates. The test statistic for `D+M` is a double mutant's residual with respect to the fitted model of GE,

$$\hat{D}_{ij}^{\mathtt{D+M}} := (\hat{Y}_{ij}^{\mathtt{D}} - \hat{Y}_{ij}^{\mathrm{pred}})/\hat{s}_{ij},$$

where $\hat{Y}_{ij}^{\mathrm{pred}}$ is the predicted double mutant phenotype (from `MoCHI`). A $p$-value is then computed for a two-sided test under the assumption that $\hat{D}_{ij}^{\mathtt{D+M}} \sim \mathcal{N}(0,1)$,

$$p_{ij}^{\mathtt{D+M}} := 2 \cdot [1 - \Phi(|\hat{D}_{ij}^{\mathtt{D+M}}|)].$$

Standard multiple testing procedures, such as Benjamini and Hochberg (1995), are used to identify significant SE pairs.

We applied `DiMSum` (using the default settings) to the simulated and empirical datasets after filtering on initial read counts. A pseudo-count of $\eta = 1$ was added to all read counts in keeping with the data processing procedures implemented in R&R. `MoCHI` (Faure and Lehner, 2024) was then used to to fit a GE model. To analyze the GB1-like simulations and the Fos-Jun DMS, we specified the form of the nonlinearity as a sum of sigmoids ("SumOfSigmoids"). For the GB1 dataset (Olson et al., 2014), we specified the nonlinearity as determined by a (1) two-state thermodynamic model ("TwoStateFractionFolded"), and a (2) three-state thermodynamic model ("ThreeStateFractionBound") in two separate analyses.

The $p$-values of R&R are bounded below by the number of bootstrap replicates, whereas those of `D+M` can, in theory, span the unit interval. Thus, to compare results from R&R with those of `D+M` on each dataset, we set the significance threshold for `D+M`, $\alpha_{DM}$, such that the false positive rate of `D+M` approximates that of R&R for a fixed $\alpha$. Namely,

$$\alpha_{DM} := \arg\min_{\alpha'} |fpr_{DM}(\alpha') - fpr_{RR}(\alpha)|, \tag{S4}$$

where $fpr_{DM}(\alpha')$ and $fpr_{RR}(\alpha)$ are the false positive rates for `D+M` and R&R evaluated at $\alpha'$ and $\alpha$, respectively.

For all analyses of the simulations, we set $\alpha = .1$. For `D+M`: GB1-like low, $\alpha_{DM} = 0.03$ and GB1-like high, $\alpha_{DM} = 0.06$. For the Fos-Jun analysis, $\alpha = .05$, and for `D+M`, $\alpha_{DM} = 3.6, 2.4$, and $3.0 \cdot 10^{-3}$. (In the Fos-Jun analysis, we computed a pseudo-false positive rate.)

#### S1.5 Two-sided $p$-value

In its application to simulated and empirical data sets, R&R exhibits asymmetry in its ability to detect positive versus negative SE in certain parts of the measurement range (see Section S2.6.8). Motivated by this observation, we propose a two-sided test.

We first estimate the distribution $\hat{D}_{(r,s)}$ (rather than its absolute value), referred to as $\tilde{F}_{(r,s)}$, and then compute,

$$p_{ij}^{\mathrm{two}} := 2 \cdot \min\left[\tilde{\mathcal{F}}_{(r,s)}(\hat{D}_{ij}), 1 - \tilde{\mathcal{F}}_{(r,s)}(\hat{D}_{ij})\right], \tag{S5}$$

where the factor of 2 accounts for multiple testing.

### S2 Supplementary results

#### S2.1 Additional simulation results

We first observe that the resulting $p$-value distribution qualitatively deviates from uniformity, with a depletion of small $p$-values (Fig. S1). The non-uniform $p$-value distribution is likely largely a consequence of overdispersion in the bootstrap sample relative to the observed data, which results from uncertainty in the estimates of the variant frequencies in the pre- and post-selection pools (Section S2.2). As a result, R&R is conservative in its detection of SE, as evidenced by low false positive rates in the majority of the measurement range (Fig. S1).

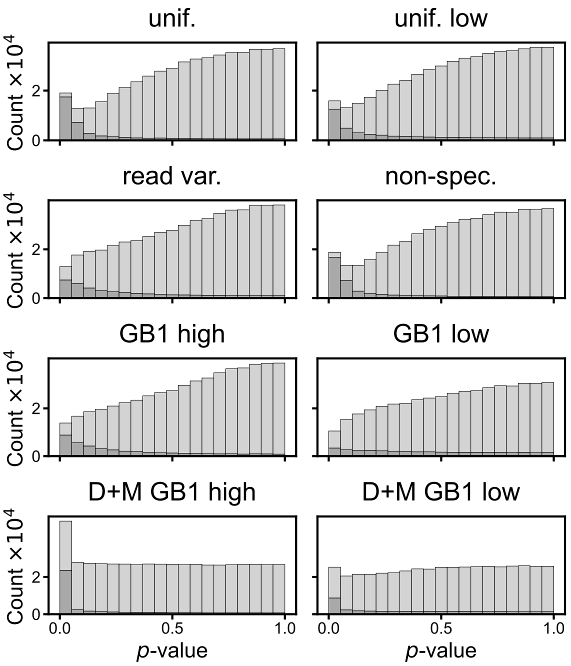

**Fig. S1. Analysis of simulated data.** Histograms of $p$-values for null mutant pairs (light gray, $\lambda_{ij} = 0$) and true interacting pairs (dark gray, $\lambda_{ij} \neq 0$) resulting from analysis of six simulation scenarios with R&R (*top three rows*). Analogous histograms for analysis of the GB1-like high and GB1-like low simulations with `D+M` (*bottom row*).

We present results for two of the simulation scenarios not detailed in the main text (see Section 5.3.3). We observe that cell frequency variation ("read variation") leads to a wholesale reduction in statistical power, while qualitatively preserving the distribution of power and false positive rates across the measurement range observed in the "uniform" simulations (Fig. S2). Non-specific binding in isolation has minimal impact on the performance of R&R (Figs. S2c and S2f). These results suggest that variation in cell counts and overall sequencing coverage are primary contributors to reductions in statistical power.

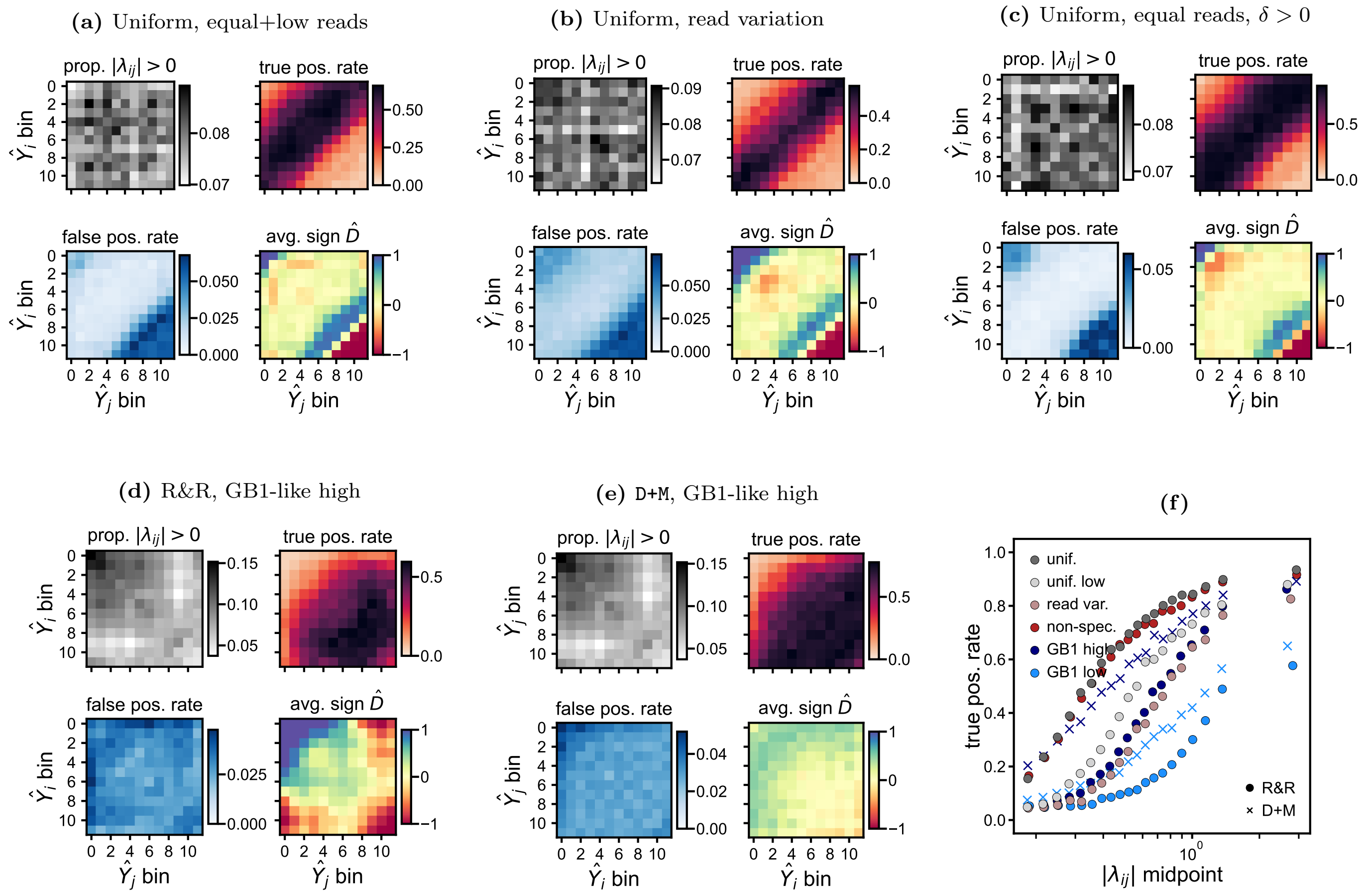

**Fig. S2. Performance of R&R on additional simulations.** R&R applied to four simulation scenarios: **(a)** Uniform single effects $\lambda_i$, equal initial counts, $C_i^0 R^0 = 10^3$ and $C_{ij}^0 R^0 = 10^2$. **(b)** Uniform $\lambda_i$ with initial cell counts sampled from the empirical GB1 read count distribution. **(c)** Uniform $\lambda_i$ with equal initial counts, $C_i^0 R^0 = 10^4$ and $C_{ij}^0 R^0 = 10^3$. **(d)** $\lambda_i$ sampled from a distribution derived from the "empirical" GB1 distribution (Otwinowski, 2018) with initial cell counts sampled as in **b**. **(e)** Analysis of the GB1-high simulations with D+M. In each **a-e**, the heatmaps represent four different quantities computed with respect to single mutant rank bins, ordered from least to most fit: The proportion of true non-zero epistatic coefficients $\lambda_{ij}$ (*top left*); the false positive rate (*bottom left*); the true positive rate (*top right*); and the average sign of the test statistics, where the average is taken over significant locus pairs (*bottom right*). Significant locus pairs are defined as those with $p$-values below $\alpha = .1$ in all cases. **(e)** True positive rates for each simulation scenario analyzed with R&R and D+M.

#### S2.2 Overdispersion in bootstrap sample.

In the simulations, the estimated fitness of a variant $i$ is given by the ratio of its pre- and post-selection reads, $\hat{N}_i^0$ and $\hat{N}_i^1$, respectively. $\hat{N}_i^0$ is Poisson distributed with mean given by the initial cell count $C_i^0$. $\hat{N}_i^1$ is Poisson distributed with mean given by $rC_i^0[p_b(1-\delta)+\delta]$, where $r$ is a global factor and $\delta$ is the probability of non-specific binding (Section 5.3.1).

When bootstrapping under the Poisson error model (Eq. 7), we assume that the pre- and post-selection reads, $\hat{N}_i^{0,b}$ and $\hat{N}_i^{1,b}$, are Poisson distributed with means given by the "observed" pre- and post-selection reads $\hat{N}_i^0 + \eta$ and $\hat{N}_i^1 + \eta$, respectively, where $\eta = 1$ is a pseudo-count. Thus, $\mathbb{E}[\hat{N}_i^{0,b}] = C^0 + \eta$ and $\mathbb{E}[\hat{N}_i^{1,b}] = rC^0(p_b(1-\delta)+\delta) + \eta$, respectively. The variance of the bootstrapped read counts, however, is inflated by a factor of two,

$$\mathbb{V}[\hat{N}_i^{0,b}] = \mathbb{E}\left[\mathbb{V}[\hat{N}_i^{0,b} + \eta | \hat{N}_i^0]\right] + \mathbb{V}\left[\mathbb{E}[\hat{N}_i^{0,b} + \eta | \hat{N}^0]\right] = \mathbb{E}[\hat{N}^0] + \mathbb{V}[\hat{N}^0 + \eta] = C^0 + C^0 = 2C^0, \tag{S6}$$

and likewise the variance of $\hat{N}_i^{1,b}$ is $2\mathbb{V}[\hat{N}_i^1]$. This result implies that the bootstrapping procedure alone is likely to produce more extreme fitness estimates than observed in the data, reducing the power of R&R to detect SE. On the other hand, the above calculation implies that the Poisson error model may capture some of the overdispersion observed in numerous sequencing datasets analyzed herein, including that of Fos-Jun (Faure et al., 2020).

#### S2.3 Additional analysis of the Fos-Jun deep mutational scan

The test-statistics and $p$-values produced by R&R were correlated across replicates (average Spearman's $\rho$ values of approximately .4 and .2, respectively, and Fig. S3), with approximately 26% of significant associations ($\alpha = 0.05$) shared among pairs of replicates.

Interactions that replicate tend to have higher double fitness values than compared to those that do not replicate (Fig. S3b). This result is consistent with the higher false positive rates observed among deleterious mutants in simulations.

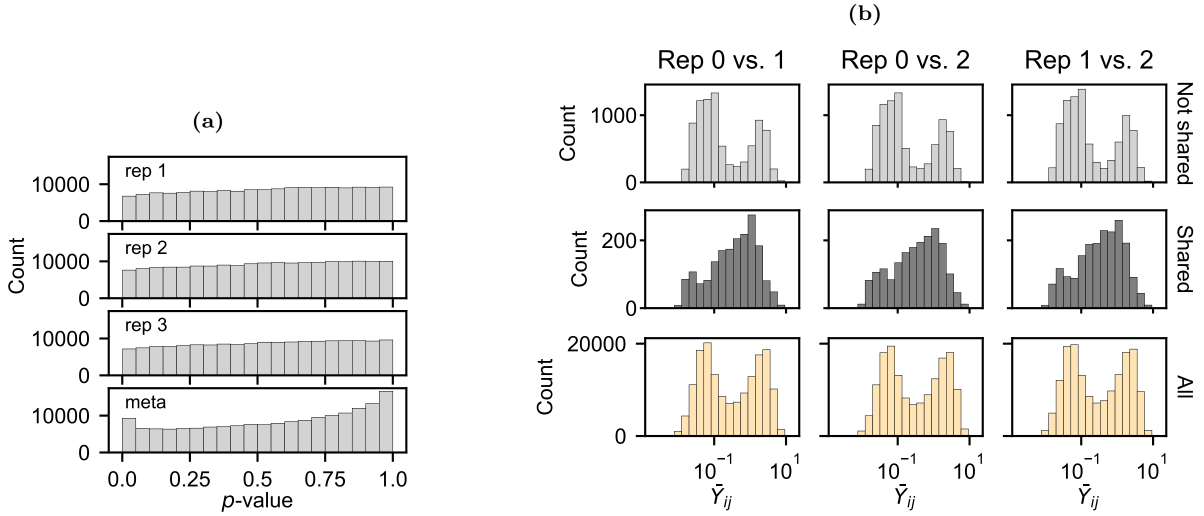

**Fig. S3. Replicability of R&R results. (a)** The $p$-value distributions for each of the three replicates (*top three rows*) and meta $p$-values computed from the three replicates (*bottom row*). **(b)** (*Top row*) Histograms of the average double mutant fitness, $\bar{Y}_{ij}$, for interactions with $p$-values below $\alpha = 0.05$ in only one of the two replicates. (*Middle row*) The same, but $\hat{Y}_{ij}$ values associated with $p$-values less than $\alpha$ in both replicates. (*Bottom row*) The same, but for all observed double mutants.

##### S2.3.1 Epistasis among protein contacts

In Section 3.2, we showed that while epistasis among enriched protein contacts is disproportionately positive, significant interactions among mutations at T8 Fos and V8 Jun are predominantly negative. Here, we demonstrate that the observed negative epistasis between T8 Fos and V8 Jun (Fig. S4) is unlikely to be explained by technical artifacts or biases in R&Rs ability to detect negative epistasis in certain parts of the measurement range (see Section 3.1 and Fig. S9).

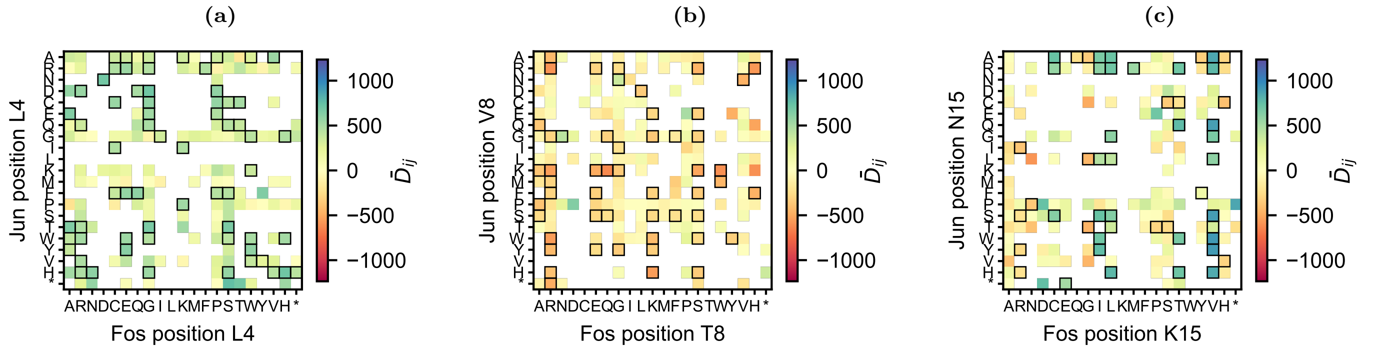

**Fig. S4. Test statistics for amino acid pairs.** The value of the test statistic $\hat{D}_{ij}$ averaged over replicates for the observed amino acid pairs for three focal position pairs **(a)** Fos L4, Jun L4; **(b)** Fos T8, Jun V8; and **(c)** Fos K15, Jun N15. Pairs of amino acids with meta $p$-values less than .05 are outlined in black.

Single mutants for both T8 Fos and V8 Jun are, on average, sequenced at higher depths in the initial sample, $\hat{N}_i^0$, than all single mutants. In addition, while we observed a weak relationship between initial single mutant reads, $\hat{N}_i^0$, and residuals, where residuals are defined as the difference between a single mutant's rank and the background-averaged rank (averaged across replicates), neither T8 Fos nor V8 Jun were atypical in this respect. All together, these observations suggest that imprecise single mutant fitness measurements did not contribute appreciably to detection of epistasis in the Fos-Jun data set

(Fig. S5). (See Section S2.5.3 for further discussion of the effect of imprecise single fitness measurements on false positive rates.) Moreover, single mutant rank and background-averaged rank were highly correlated in each replicate (Spearman's $\bar{\rho} \approx .99$ averaged across replicates, and see Fig. 5b).

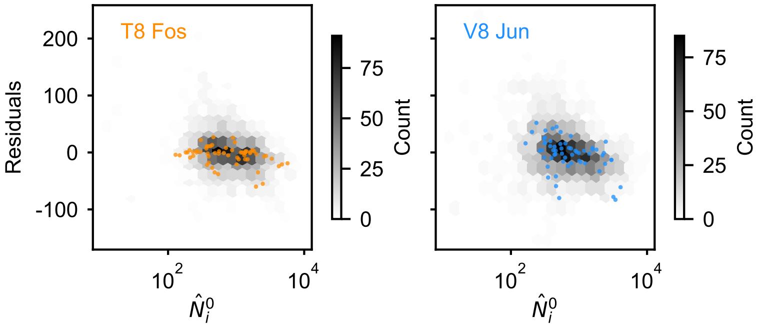

**Fig. S5. Joint distribution of single mutant read depth and residuals.** Two dimensional histograms of the average single mutant read depth and residuals, defined as the difference between a single mutant's rank and its background-averaged rank, averaged across replicates, for Fos single mutants (left) and Jun single mutants (right). Mutations of T8 Fos and V8 Jun are shown in orange and blue on the left and right, respectively.

In addition, the signal of predominant negative epistasis persists when the data are analyzed with a two-sided test (Eq. S5) and with `D+M`, which both exhibit less sign bias in detection of epistatic effects throughout the measurement range (Fig. S6 and see Section S2.3.2). To quantify whether the distribution of signed-$D$ statistics could have arisen by chance, we assume that the number of positive $D$-statistics in a given pair of rank bins is binomially distributed with probability given by the observed fraction of significant positive $D$-statistics across all positions. Letting $s_{ab}$, $n_{ab}$, and $\hat{p}_{ab}$ denote the number of significant positive $D$-statistics, the number of significant $D$-statistics, and the fraction of significant $D$-statistics across all positions, in the bin pair $(a, b)$, the pseudo-log-likelihood is given by,

$$\ell\ell(\mathbf{s}, \mathbf{n}) \coloneqq \sum_{a,b} \log \left( \text{Binomial}(s_{ab}; n_{ab}, \hat{p}_{ab}) \right), \tag{S7}$$

where the sum is over all bin pairs $(a, b)$; and Binomial$(\cdot; \cdot, \cdot)$, denotes the binomial likelihood. We compute the pseudo-likelihood for each replicate separately and procedure (R&R "one" and two-sided $p$-values and `D+M`). To evaluate significance (in each replicate), we compute the likelihoods of a $n = 1000$ random samples from the likelihood model. In each analysis, two of the three $p$-values were below .1, with maximum $p$-values of .18, .17, and .21, respectively, providing evidence in favor of our claim of strong negative epistasis between T8 Fos and V8 Jun.

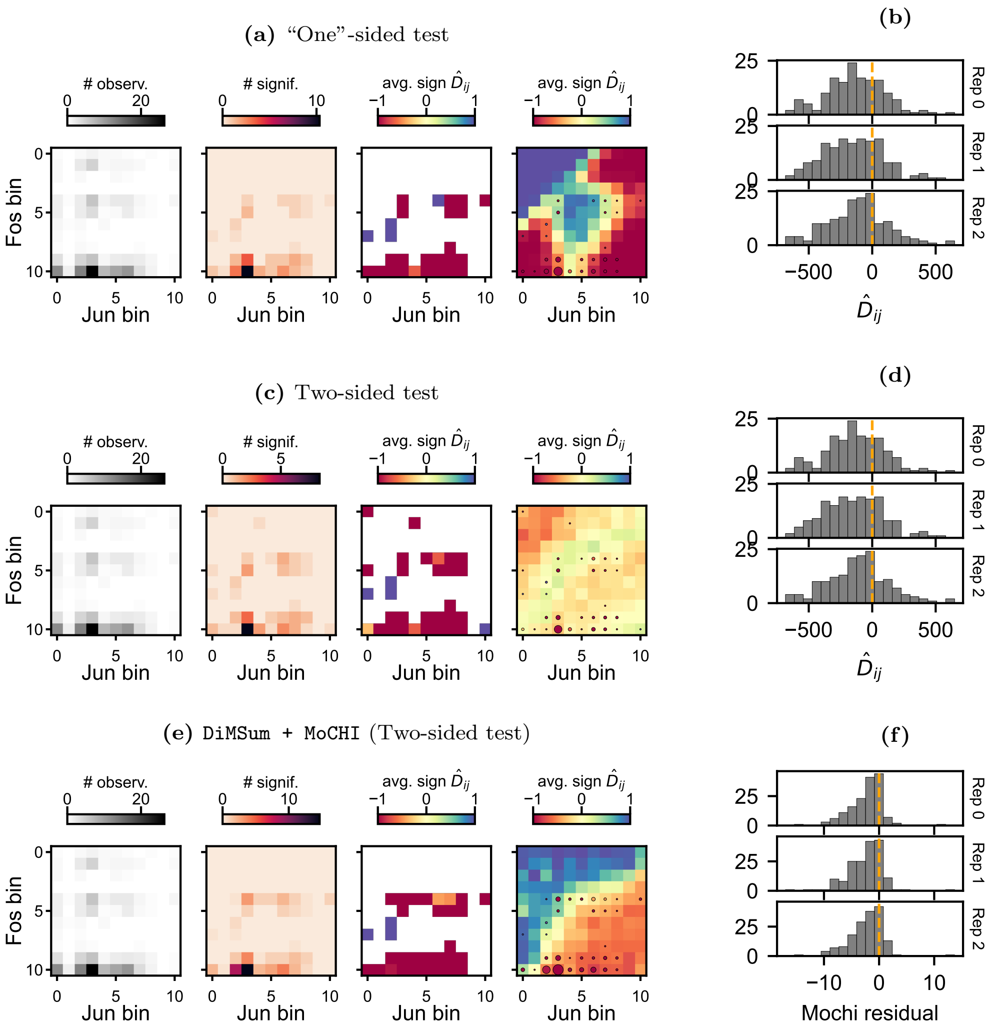

**Fig. S6. Negative epistasis among T8 Fos and V8 Jun mutations.** **(a)** Fos ($y$-axis) and Jun ($x$-axis) mutations were sorted into bins based on their single fitness estimates, from lowest to highest. The number of observed double mutants (first column), the number of interactions with $p$-values less than $\alpha$ (second column), and the average sign of such interactions (third column), among T8 Fos and V8 Jun mutant pairs for the one-sided test ($\alpha = .05$), averaged across the three replicates. In the rightmost column, the average sign of interactions with $p$-values below $\alpha$ among all mutant pairs overlaid with points denoting the average sign of all such interactions among T8 Fos and V8 Jun (the quantity displayed in the third column). The size of the point corresponds to the number of $p$-values below $\alpha$ for T8 Fos and V8 Jun. **(b)** Histograms of all $\hat{D}_{ij}$ values for T8 Fos and V8 Jun amino acid combinations in each replicate. **(c)** The same as in **a**, but for the two-sided test with $\alpha = .048$ in each replicate. **(d)** The same as **(b)**. **(e)** The same as in **a**, but for `D+M` with $\alpha \approx 3 \cdot 10^{-3}$ in each replicate. **(f)** Histograms of the normalized residuals from `D+M` for each replicate.

We do not, however, exclude the possibility that the overwhelming positive epistasis observed among protein contacts is a consequence of R&Rs (and `D+M`s) inability to detect negative epistasis among deleterious mutations. For example, the result of positive epistasis among L4 Fos and L4 Jun mutations is unremarkable with respect to the detection biases of R&R and `D+M` (Fig. S7). However, the positive epistasis persists, and is unlikely, under a two-sided test performed with R&R (Fig. S7c,d). At the time, substantial negative epistasis is observed among some enriched contact pairs, such as K15 Fos and N15 Jun (Fig. S4).

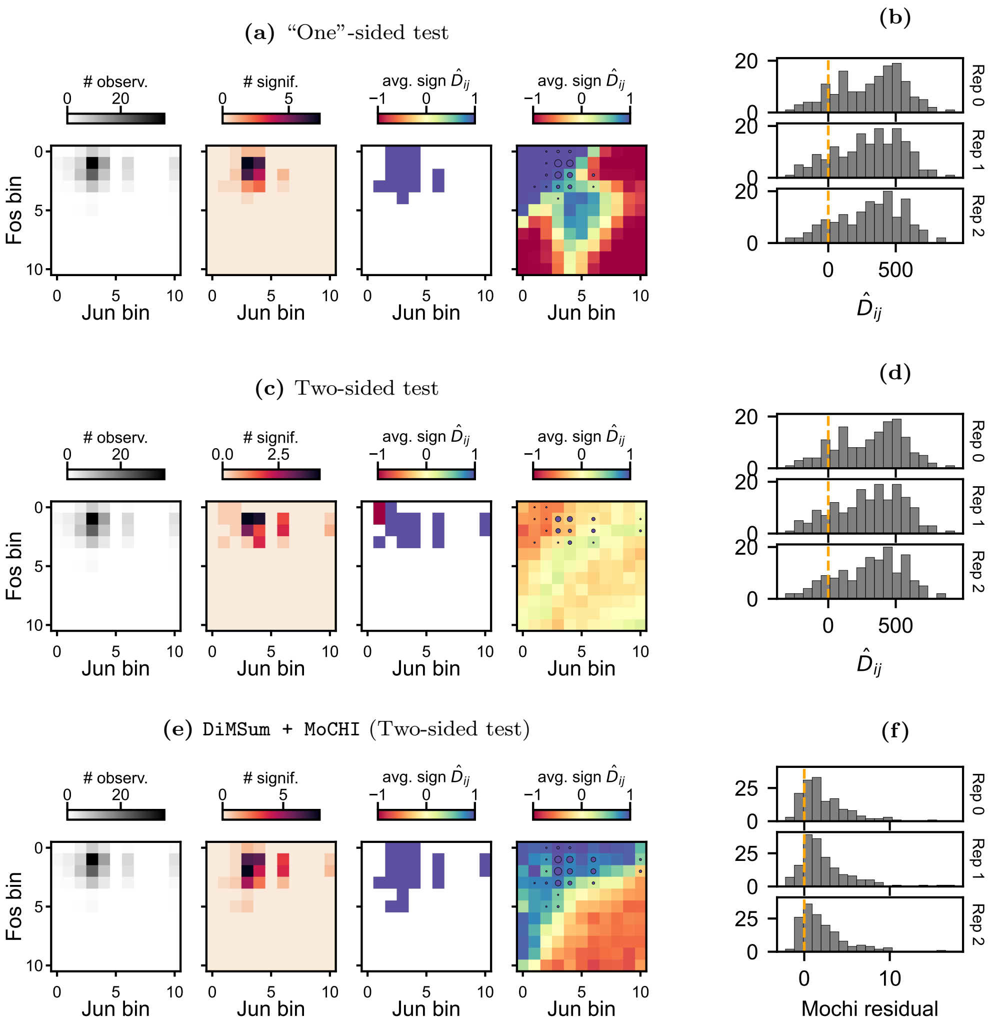

**Fig. S7. Positive epistasis among L4 Fos and L4 Jun mutations.** The same as in Fig. S6 but for L4 Fos and L4 Jun.

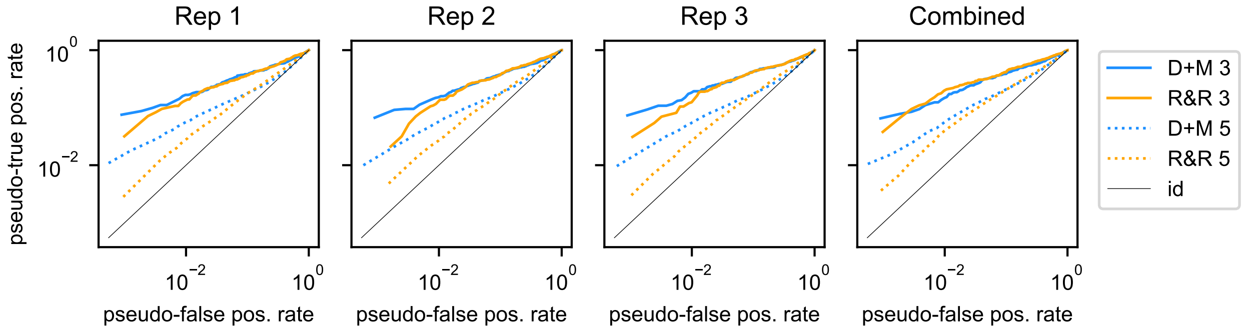

**Fig. S8. Receiver operating curves for the analysis of the Fos-Jun dataset**. Pseudo-true positive rate as a function of pseudo-false positive rates for R&R (orange) and `D+M` (blue) computed with respect to true interactions defined as amino acids within 3Å(solid lines) and 5Å(dotted lines) in each replicate (first three columns) and the combined analysis (rightmost column).

##### S2.3.2 Comparison to DiMSum and MoCHI

To detect SE with `DiMSum` (Faure et al., 2020) and `MoCHI` (Faure and Lehner, 2024), referred to as `D+M`, we followed the procedure described in Section S1.4. We compared the methods in terms of their *pseudo*-accuracy and recall, where *pseudo*-true interactions were defined here as pairs within 5Å in the crystal structure, at the level of (1) amino acids and (2) position pairs.

**Accuracy at the amino acid level.** To compare R&R and D+M at the amino acid level, we fixed the D+M $\alpha$ threshold such that the pseudo-false positive rate was the same for both methods in each replicate. D+M exhibited similar, though less pronounced, bias in the sign of detected epistasis throughout the measurement range (Fig. S9d-f). In addition, D+M was relatively less powered to detect SE among deleterious mutations, but exhibited lower pseudo-false positive rates among these pairs (Fig. S9d-f and Fig. S10d-f).

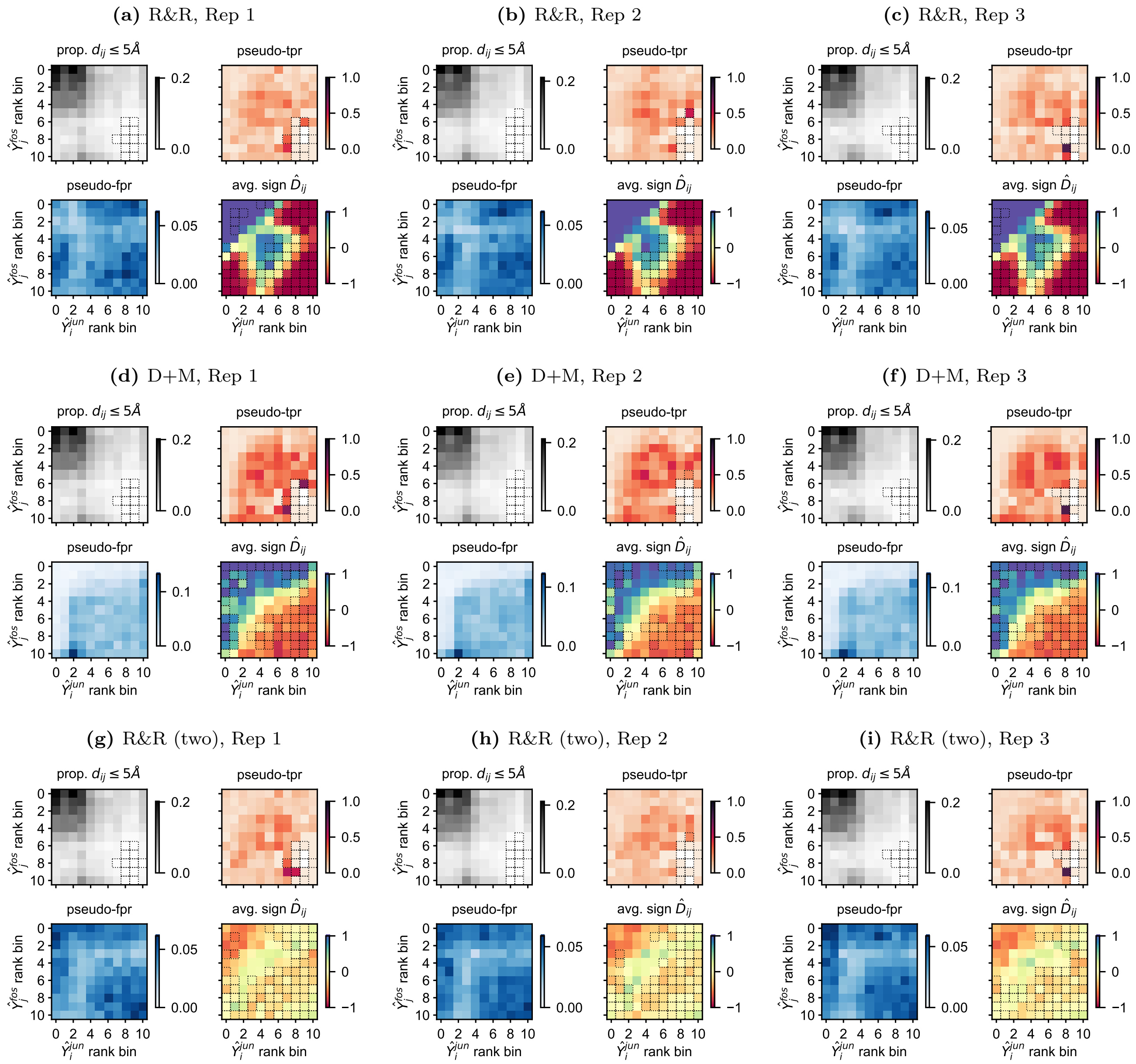

**Fig. S9. Biases in the detection of specific epistasis in the Fos-Jun dataset.** In each heatmap, single mutants are sorted into bins based on their single mutant ranks, with Jun mutations on the $x$-axis and Fos mutations on the $y$-axis. In the top left of each subplot, each square represents the proportion of mutant pairs that correspond to protein contacts, defined as amino acids within 5Å. For R&Rs "one"-sided test (**a-c**), D+M (**d-f**), and R&Rs two-sided test (**g-i**), the pseudo-false (bottom left) and -true positive (top right) rates are shown for all pairs of rank bins. For R&R, the significance threshold was set to $\alpha = .05$ across replicates. For D+M, $\alpha = 3.6, 2.4, 3.0 \cdot 10^{-3}$ in each replicate, respectively, to match the pseudo-false positive rate of R&R. For R&Rs two-sided test, $\alpha-$ The average sign of the test statistic, across significant pairs of mutants within each bin, is shown for each pair of rank bins (bottom right). In each subplot, squares are outlined when fewer than five measurements contribute to the estimate.

To further compare the two methods, we computed a receiver operating characteristic curve, or ROC curve, for each method applied to each replicate separately. In addition, we computed ROC curves for D+M applied to the fitness estimates combined across replicates, as well as, the meta $p$-values estimated from the R&R results from the three replicates. To estimate the curve, we computed the pseudo-true and false positive rates for 1000 evenly spaced quantiles, separately for each method and replicate (Fig. S8). We observe that in both methods the pseudo-true positive rate increases faster than the pseudo-false negative rate, i.e., the ROC curves fall above the identity line, with D+M outperforming R&R when amino acids within 5Å in the crystal structure are specified as true interacting pairs (dotted lines). When the distance is reduced to 3Å, the methods are more comparable. Assuming that more proximal amino acids are more likely to exhibit strong interactions, this result is consistent with D+M being better powered to detect larger effect SE in simulations.

**Accuracy at the position pair level.** We test whether particular pairs of positions are enriched for SE, described in Section 5.2.5, using the $p$-values from both R&R and D+M. (Though, here, we do not remove null alleles.) We consider the pseudo-accuracy and recall of enrichment tests with respect to contacts within 5Å over a range of $\alpha$ values. We find that R&R and D+M perform similarly across a range of $\alpha$ values when the test is conducted within replicates or when we compare the R&R meta $p$-values with D+M applied to fitness values estimated from the three replicates simultaneously (Fig. S11).

We leave a detailed comparison of methods for detecting SE to future work. Until the biases of SE detection methods, including D+M and other linear and non-linear regression procedures, are understood, we encourage practitioners to apply several procedures, including R&R, to analyze their data.

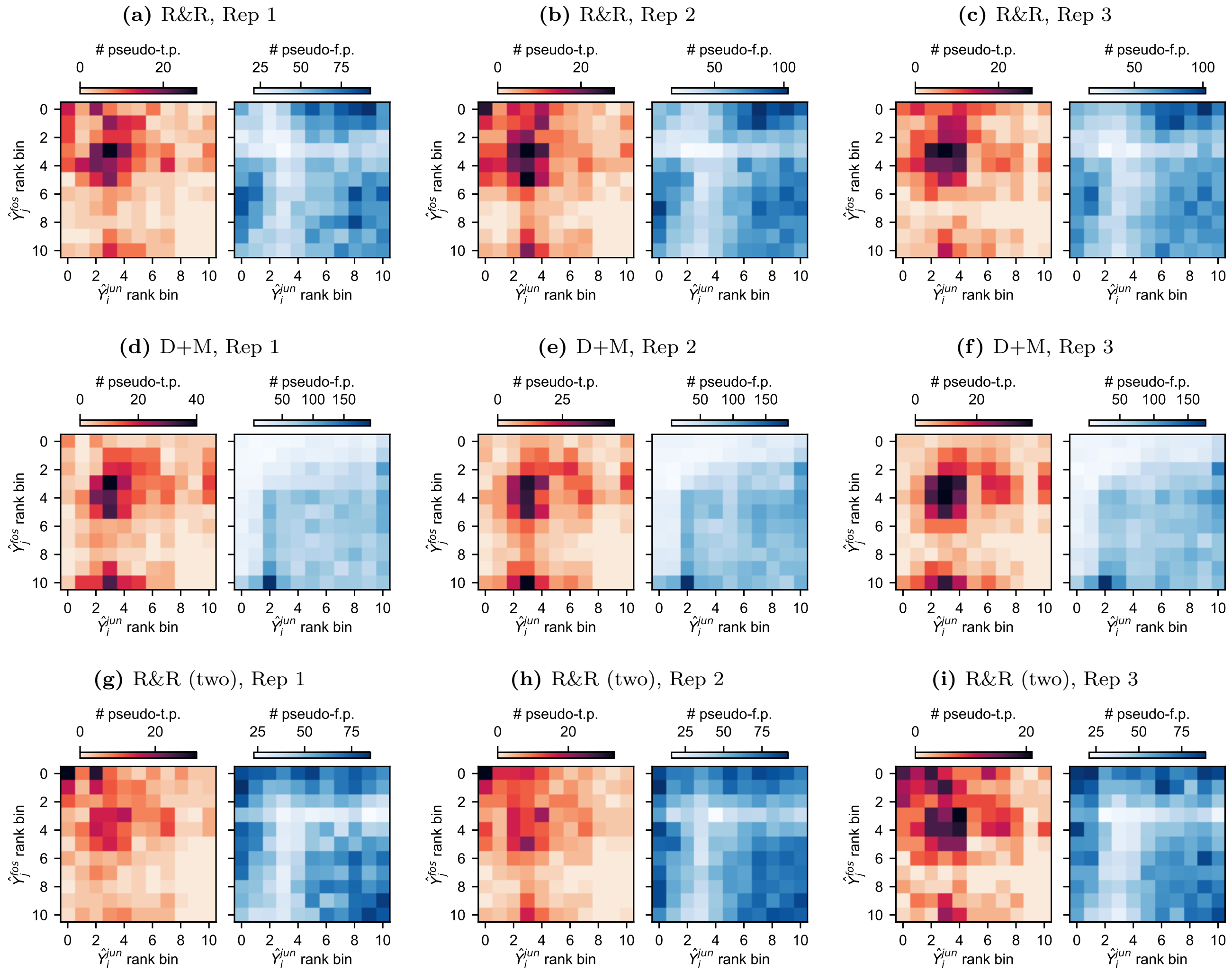

**Fig. S10. Pseudo-true and false positives in the Fos-Jun dataset. (a-c)** In each heatmap, single mutants are sorted into bins based on their single mutant ranks, with Jun mutations on the $x$-axis and Fos mutations on the $y$-axis. The absolute numbers of pseudo-true (left) and false (right) positives in the analysis of the Fos-Jun DMS with R&R ($\alpha = .05$) are plotted for each replicate. **(d-f)** The same quantities estimated from analysis with D+M, where the significance threshold was chosen for parity with the R&R analysis ($\alpha = 3.6, 2.4, 3.0 \cdot 10^{-3}$, respectively). And likewise, **(g-i)** for the two-sided analysis with R&R.

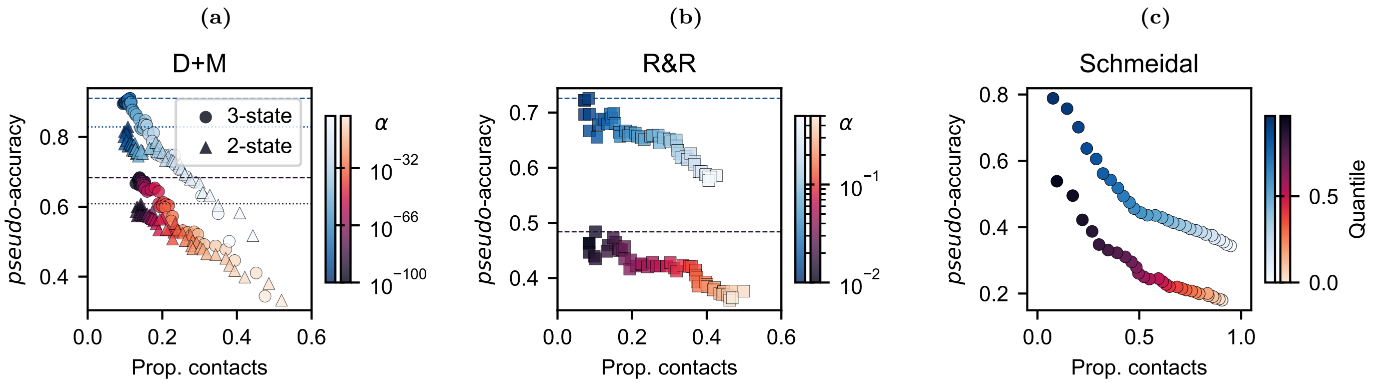

**Fig. S12. Contact prediction accuracy in GB1 using two- and three-state models. (a)** The proportion of enriched pairs which correspond to amino acids within 5Å (pink) and 8Å (blue) plotted as a function of the proportion of total contacts (at each distance threshold) identified in two- (triangles) and three-state (circles) models fitted with `D+M`. Each point represents a different threshold $\alpha$ used in the enrichment test, with color denoting $\alpha$. A position pair was identified as a predicted contact if its enrichment $p$-value exceeded a false discovery rate of $\alpha_{BH} = 10^{-2}$ (Benjamini and Hochberg, 1995). **(b)** The same as in **a**, but for R&R. **(c)** The same as in **a**, except instead of signifiance thresholds, each point corresponds to a quantile, below which a position pair is said to be "significant".

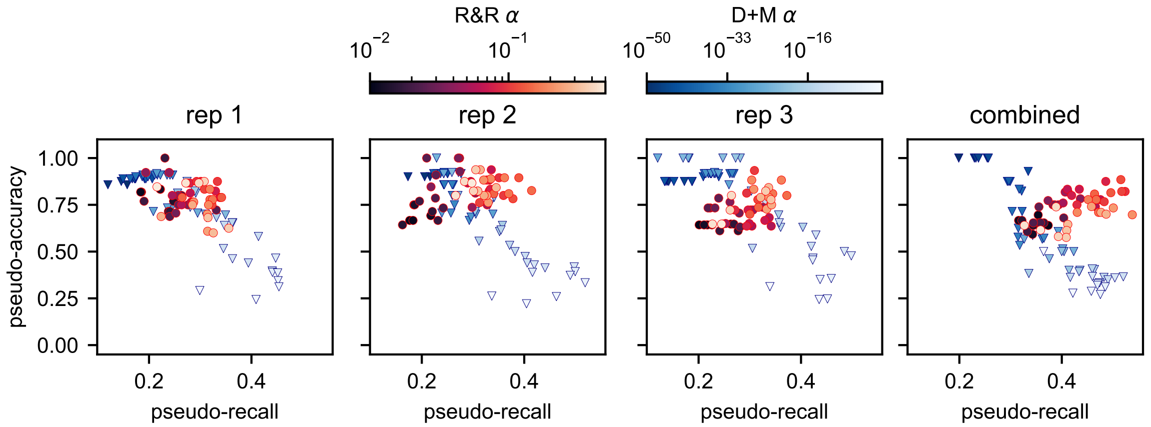

**Fig. S11. Contact prediction accuracy and recall in Fos-Jun for R&R and `D+M`.** Pseudo-accuracy as a function of pseudo-recall for a range of $p$-value thresholds $\alpha$ for R&R (red circles) and `D+M` (blue triangles). The color of the point denotes $\alpha$. Pseudo-accuracy is defined as the number of enriched position pairs within 5Å divided by the total number of enriched pairs after correcting for multiple testing ($\alpha_{BH} = 0.05$). Pseudo-recall is the proportion of total contacts detected.

#### S2.4 Contact accuracy in GB1

We estimated contact prediction accuracy for two- and three-state models fitted with `D+M` for various significance thresholds $\alpha$, where predicted contacts exceeded a false-discovery rate of $\alpha_{BH} = 10^{-2}$ (Fig. S12 and see below for a description of the the three-state model). The three-state model improved pseudo-contact accuracy from 61% to 68% when contacts were identified as amino acids within 5Å, and from 83% to 91% at 8Å (pink and blue points in Fig. S12, respectively). R&R achieved maximum prediction accuracies of 48% and 73% across different $\alpha$ values, but recovered approximately twice as many contacts as either the two- or three-state `D+M` models at their respective accuracy maxima (Fig. S12). Schmiedel and Lehner (2019), which implemented a heuristic procedure for identifying interactions, achieved high *pseudo*-accuracy when only the "top" pairs were retained.

##### S2.4.1 Global epistasis in higher dimensions

R&R relies on the assumption of a single underlying latent trait $\Lambda$. We refer to this scenario as single trait GE. In practice, however, the latent space may be higher-dimensional. For example, a three-state model, with two underlying latent traits, is thought to provide a better description of the DMS of GB1 (Otwinowski, 2018)—an example we consider in Section 3.3.

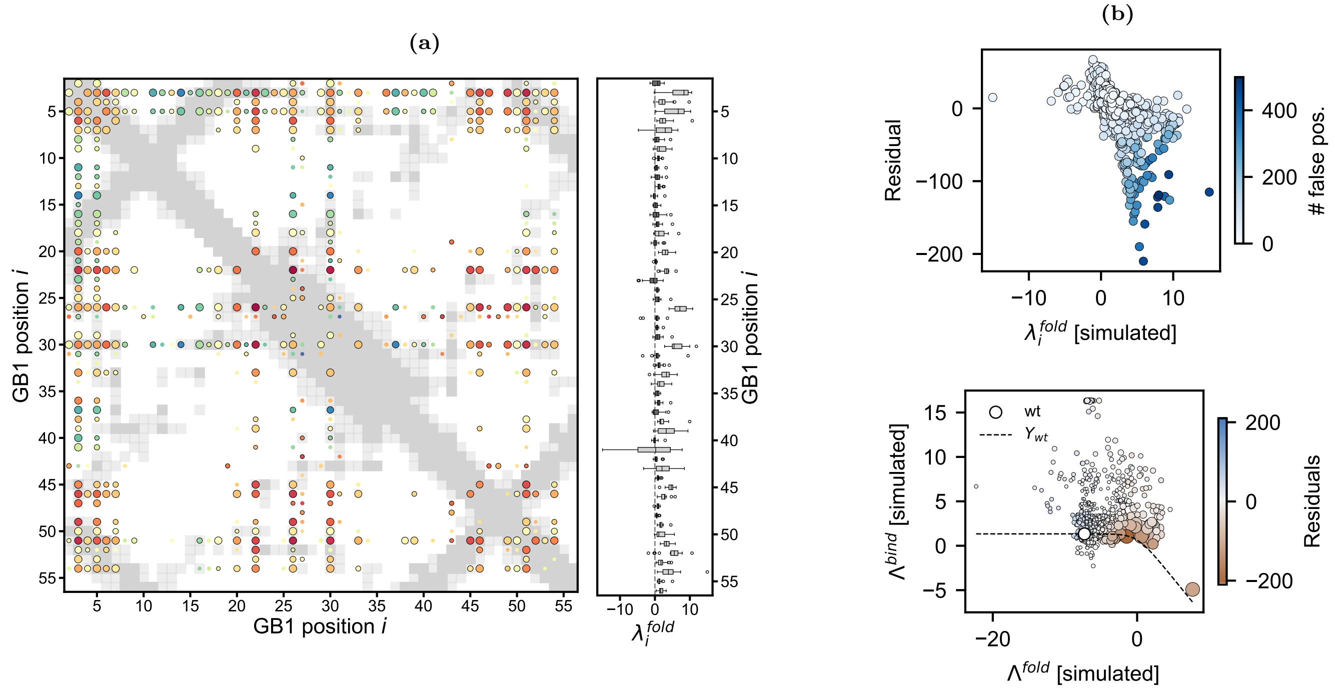

**Fig. S13. Simulations under a three-state thermodynamic model.** A DMS was simulated under the assumption of a three-state thermodynamic model with folding and binding energies estimated in Otwinowski (2018) and no specific epistasis (SE). **(a)** Position pairs enriched for SE ($\alpha_{BH} = .1$) indicated by points colored by the average sign of SE among significant amino acid pairs. The dark gray and light gray boxes indicate position pairs within 5Å and 8Å, respectively. **(b)** Box plots of the folding energies, $\lambda_i^{\text{fold}}$, taken from (Otwinowski, 2018) for each position. Light gray indicates that the average $\lambda_i^{\text{fold}}$ significantly deviates from the bulk average after a multiple testing correction. **(c)** Residuals as a function of $\lambda_i^{\text{fold}}$ (*top*). In the *bottom* of **c**, each mutant is plotted in the two-dimensional latent space defined by the folding and binding energies, $\Lambda^{\text{fold}}$ and $\Lambda^{\text{bind}}$. The size of the point corresponds to the number of false positives associated with the mutant, and the color to its residual.

In a three-state model, the protein may occupy three distinct states, here: unfolded and unbound ($uu$), folded and unbound ($fu$), and folded and bound ($fb$), as in Otwinowski (2018),

$$uu \xrightleftharpoons{\Lambda^{\text{fold}}} fu \xrightleftharpoons{\Lambda^{\text{bind}}} fb,$$

where $\Lambda^{\text{fold}}$ and $\Lambda^{\text{bind}}$ are the folding and binding energies of the protein, respectively. (The three states could also pertain to different protein conformations.) Under this model, the phenotype of a variant $i$ is determined by the relative energy of the $fb$ state,

$$Y_i \propto \frac{e^{-(\Lambda_i^{\text{bind}}+\Lambda_i^{\text{fold}})}}{1+e^{-\Lambda_i^{\text{bind}}}+e^{-(\Lambda_i^{\text{bind}}+\Lambda_i^{\text{fold}})}} = \left[e^{\Lambda_i^{\text{bind}}}(e^{\Lambda_i^{\text{fold}}}+1)+1\right]^{-1}, \tag{S8}$$

and, the *partial* derivatives of $Y_i$ with respect to the folding and binding energies are given by,

$$\left.\frac{\partial Y_i}{\partial \Lambda_i^{\text{fold}}}\right|_{\Lambda_i^{\text{bind}}} = -Y_i^2 e^{\Lambda_i^{\text{bind}}} e^{\Lambda_i^{\text{fold}}} < 0 \quad \forall \Lambda_i^{\text{fold}} \qquad \text{and} \qquad \left.\frac{\partial Y_i}{\partial \Lambda_i^{\text{bind}}}\right|_{\Lambda_i^{\text{fold}}} = -Y_i^2 e^{\Lambda_i^{\text{fold}}} e^{\Lambda_i^{\text{bind}}} < 0 \quad \forall \Lambda_i^{\text{bind}}. \tag{S9}$$

In other words, $Y_i$ is monotonic when either the folding or binding energies is held fixed. However, the ordering of two mutations now depends on *both* their folding and binding energies:

$$\begin{aligned} \{Y_i > Y_j\} &\iff e^{\Lambda_i^{\text{bind}}}(e^{\Lambda_i^{\text{fold}}}+1) > e^{\Lambda_j^{\text{bind}}}(e^{\Lambda_j^{\text{fold}}}+1) \\ &\iff e^{\Lambda_i^{\text{bind}}-\Lambda_j^{\text{bind}}} > \frac{e^{\Lambda_j^{\text{fold}}}+1}{e^{\Lambda_i^{\text{fold}}}+1}. \end{aligned} \tag{S10}$$

Therefore, when both mutants are very stable, $\Lambda_j^{\text{fold}}, \Lambda_j^{\text{fold}} \ll 0$, the right hand side of Eq. S10 is approximately one, and the binding energies of the mutants determine their pairwise ordering. In this regime, the three-state model is consistent with single-trait GE. (This condition is given as $\Lambda^{\text{fold}} < 0 < \Lambda^{\text{bind}} \approx \Lambda$ in Otwinowski (2018).) However, when the mutants are unstable, $\Lambda_j^{\text{fold}} > 0$, the pairwise ordering depends on both their folding and binding energies, and single-trait GE no longer holds.

**Simulations.** We apply R&R to data simulated under the assumption of a three-state model, with the estimated single mutant effects on $\Lambda_i^{\text{bind}}$ and $\Lambda_i^{\text{fold}}$ are specified by their estimates in Otwinowski (2018). We do not include SE, but otherwise follow the simulation procedures described in Section 5.3.1 for the *Equal cell counts* simulation.

The three-state simulations qualitatively reproduce the results of the empirical analysis, with spurious SE inferred for position pairs with above average folding energies (Fig. S13). In addition, we observe an approximately linear relationship between a mutant's residual and its folding energy, $\lambda_i^{\text{fold}}$, as well as, higher false positives for marginally stable mutants (Fig. S13b).

#### S2.5 Additional analysis of the PDZ3-CRIPT deep mutational scan

##### S2.5.1 Epistasis among protein contacts

Enrichment analysis in PDZ3-CRIPT recovered fewer protein-ligand contacts than in other data sets (Fig. S14). This result may reflect a more sparse epistatic map with longer range interactions, but may also reflect technical artifacts. While *pseudo*-accuracy consistently declined with increase $\alpha$, the proportion of identified contacts did not (compare Fig. S14b to Fig. S12b).

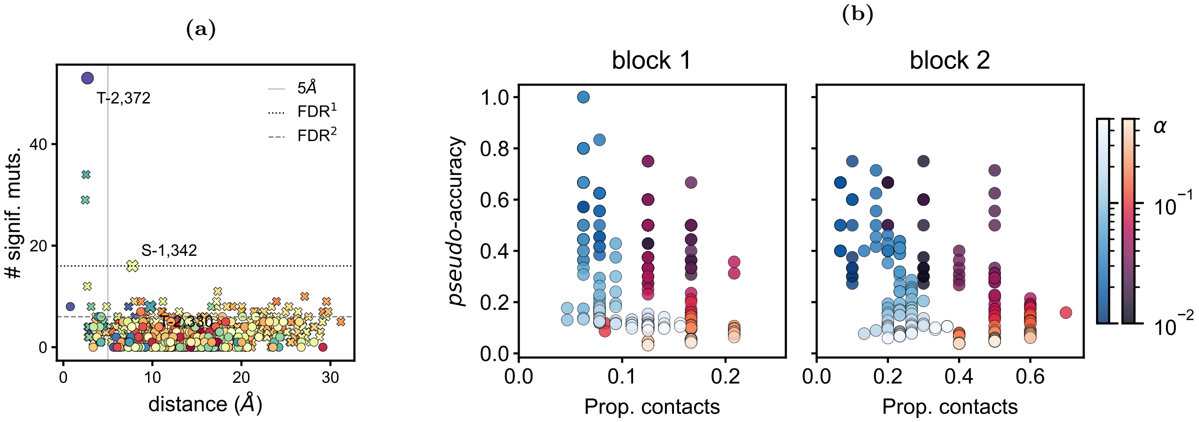

**Fig. S14. Contact prediction accuracy in PDZ3-CRIPT. (a)** The number of significant amino acid combinations for CRIPT and PDZ3 position pairs ($\alpha_1 = .034$, $\alpha_2 = .021$ for blocks 1 and 2, respectively) as a function of physical distance in the crystal structure (Å). Each point is colored by the average sign of the significant amino acid pairs. The dotted and dashed lines denote the number of significant pairs required to achieve significance after multiple testing (FDR, $\alpha = 10^{-2}$), for blocks 1 and 2, respectively. The vertical gray line denotes 5Å. **(b)** The same quantities plotted in Fig. S12a, but for blocks 1 (left) and 2 (right) of PDZ3-CRIPT with a false discovery rate of $\alpha_{BH} = .1$. Here, the input $p$-values are the meta-$p$-values computed using Fisher's method.

We provide a microscopic view of epistasis among four of the top enriched position pairs in (Fig. S15).

##### S2.5.2 Batch effects

In Faure and Lehner (2024), the first and latter 50 amino acids of the 100 amino acid region of PDZ3 (303-403aa) were processed in separate batches. Read counts in the second block (353-396aa after filtering, see Section 5.1.3) were consistently higher than in the first block (303-342aa after filtering) for both single and double mutants (Fig. S16b,d). When the data sets were combined, variants in block 2 tended to have lower single and double mutant ranks than those of block 1 (Fig. S16a,c).

The higher average single and double mutant ranks for batch 1 are likely, in part, a consequence of lower read depth. Fitness estimates derived from lower initial read counts, $\hat{N}_i^0$, have greater variance, and are more apt to exhibit more extreme values. Consider a mutant with for which $\hat{N}_i^0 = 10$ and $\hat{N}_i^1 \sim \text{Poisson}(10)$, i.e., $Y_i = 1$. The probability that $\hat{Y}_i$ exceeds $Y_i$ by 50% is non-negligible at approximately 0.08. When we increase the read (and cell) count by a factor of 10, the probability of such grave mis-estimation is less than $10^{-5}$. Therefore, in a data set with heterogeneous initial read counts, and thus cell count, distributions, we expect larger errors in the ranking of low read count variants.

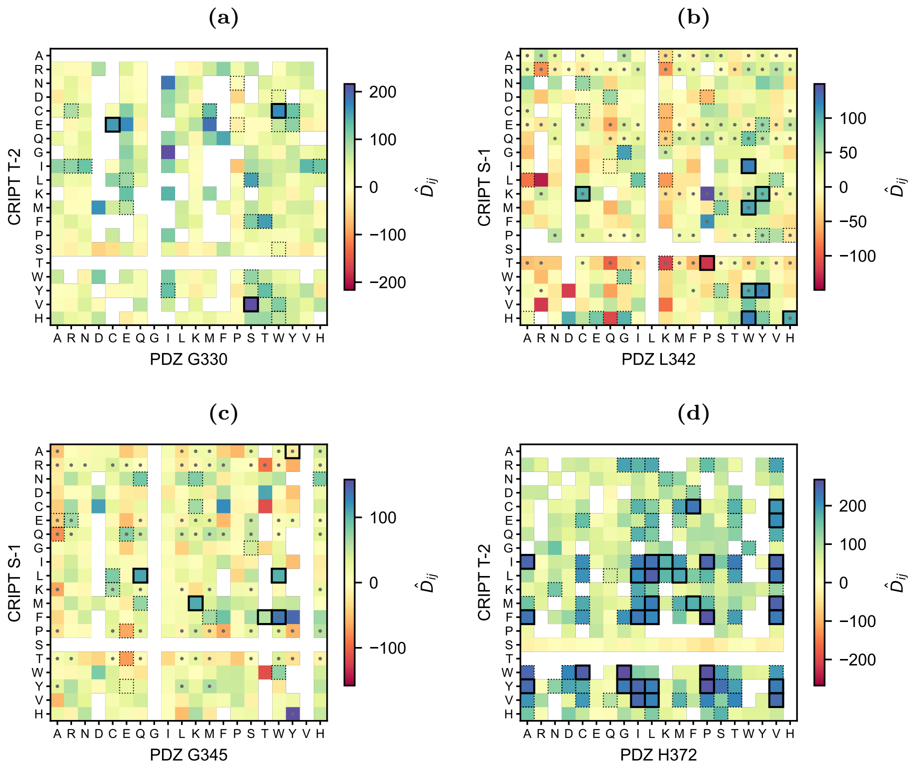

**Fig. S15. Amino acid interactions among focal position pairs.** Each heatmap shows the value of the test statistic for each combination of amino acids for a given position pair. Test statistics with $p$-values below $\alpha = 10^{-2}$ and $10^{-1}$ are outlined in solid and dotted boxes, respectively. Replicate-averaged double mutant phenotype values equal to or exceeding the wildtype fitness are indicated with gray points. **(a)** CRIPT T-2 and PDZ3 G330; **(b)** CRIPT S-1 and PDZ3 L342; **(c)** CRIPT S-1 and PDZ3 G345; and, **(d)** CRIPT T-2 and PDZ3 H372.

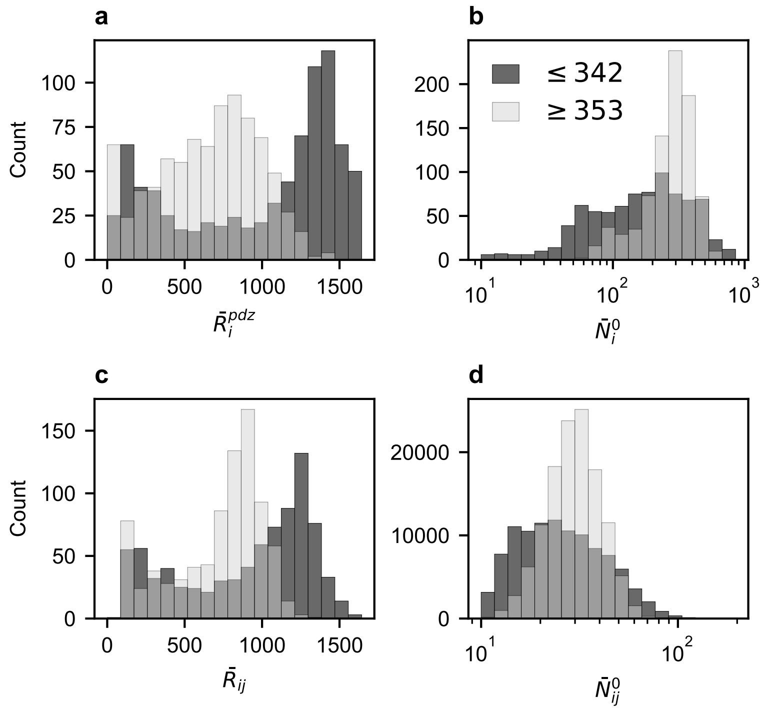

**Fig. S16. Batch effects in the PDZ3-CRIPT data set.** Histograms of **(a)** single mutant ranks and **(b)** double mutant ranks, with block 1 variants (303-342aa) in dark gray and block 2 variants (353-396aa) in light gray. Histograms of initial read counts, averaged over replicates, for **(b)** single and **(d)** double mutants.

##### S2.5.3 Comparison of R&R results across replicates

The $p$-value distributions were approximately uniform in each replicate, with qualitative depletion of small $p$-values (Fig. S17). The test-statistics and $p$-values computed from the PDZ3-CRIPT dataset were much less correlated across replicates than those of Fos-Jun, $\bar{\rho}_p = 0.05, 0.02$ and $\bar{\rho}_{\hat{D}} = 0.12, 0.12$, in blocks 1 and 2, respectively.

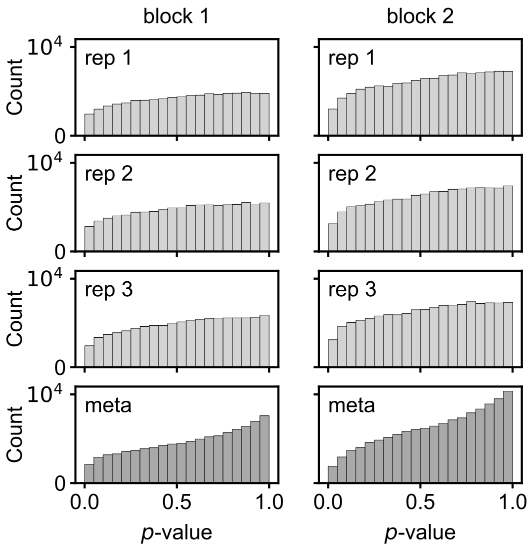

**Fig. S17. Distributions of $p$-values for PDZ3-CRIPT.** (*Top 3 rows*) Histograms of the $p$-values from each replicate for blocks 1 (left column) and 2 (right column). (*Bottom row*) Meta-$p$-values computed from the three replicates for each block.)

We suspect that lower sequencing depth—particularly of the single mutants—contributed to lower replicability in the PDZ3-CRIPT dataset. The average number of initial read counts for PDZ3 single mutants ($\bar{N}_i^0 \approx 220$ and 287) was an order of magnitude lower than for Fos and Jun mutants ($\bar{N}_i^0 \approx 1{,}022$ and 931 for Fos and Jun, respectively) and two orders of magnitude lower than for CRIPT mutants ($\bar{N}_i^0 \approx 7{,}816$ and 6,332).

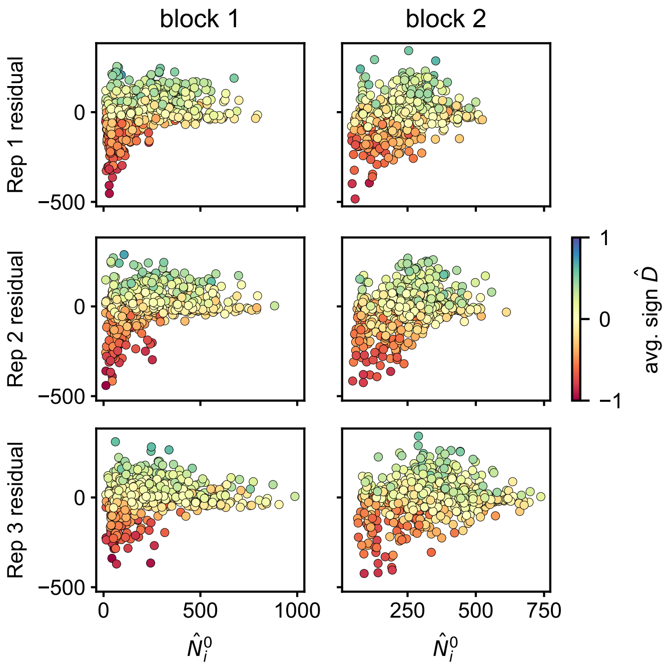

**Fig. S18. Imprecise single fitness measurements.** Residuals plotted as a function of initial single mutant sequencing depth, $\hat{N}_i^0$, for each replicate (row) and block (column). Each point is colored by its average test statistic value $\hat{D}_{ij}$ across all $j \neq i$.

As described above, low initial read counts may result in overestimation of fitness. Indeed, we observe that mutants with large negative residuals tend to have lower initial read counts (Fig. S18). Noisier fitness estimates for PDZ3 single mutants likely contributed to a reduced Spearman's correlation between single mutant rank and "average" rank across all backgrounds. In particular, "average" rank exhibited much more variation among the fitter 50% of variants (Fig. 5c). Together, these observations imply (1) reduced power to detect epistatic effects among "fit" variants and (2) pervasive bias due to low precision single fitness measurements.

#### S2.6 Theory

In this section, we provide theoretical justification for (1) privileging rank statistics as a framework for GE and (2) detecting SE in the presence of GE with R&R.

##### S2.6.1 Kendall's tau.

To build intuition, we begin our investigation with an alternative measure of rank correlation, Kendall's $\tau$. This statistic relies only on pairwise order statistics and is therefore analytically more tractable. The estimate of $\tau_{ij}$ for two mutations $i$ and $j$ is defined as,

$$\hat{\tau}_{ij} := \frac{2}{|\mathcal{A}_{ij}|(|\mathcal{A}_{ij}|-1)} \sum_{\substack{n<m \\ n,m\in A_{ij}}} \mathrm{sgn}(\hat{Y}_{im}-\hat{Y}_{in})\mathrm{sgn}(\hat{Y}_{jm}-\hat{Y}_{jn}) \tag{S11}$$

where $\mathrm{sgn}(x)=1$ for $x>0$ and $\mathrm{sgn}(x)=-1$ for $x<0$; $\mathcal{A}_{ij}$ is the set of all mutants observed in both backgrounds $i$ and $j$, and $|\mathcal{A}_{ij}|$ the size of this set. For succinctness, we have suppressed the amino acid indexing in the subscript.

In practice, ties may occur. In this case, $\mathrm{sgn}(0)=0$ (In expectation, this is analogous to sampling a signed Bernoulli random variable with equal probability.)

##### S2.6.2 Probability of correct pairwise ordering

Under the assumption of GE (Eq. 1) and when $\frac{dg}{d\Lambda}>0$, $\lambda_m>\lambda_n \iff Y_{im}>Y_{in}$. However, the probability that the measured values also exhibit this relation, i.e., $\hat{Y}_{im}>\hat{Y}_{in}$, depends on (1) the slope of $g$ in the neighborhood of $\Lambda_i$; (2) the measurement precision; and (3) the magnitude of the difference of the effects, $|\lambda_m-\lambda_n|$. To illustrate, we consider the probability that the measured phenotypes are correctly ordered, for $\lambda_m$ and $\lambda_n$ small,

$$P^i_{mn} := \mathbb{P}\{\hat{Y}_{im}>\hat{Y}_{in}\} \approx \mathbb{P}\left\{g'(\lambda_i)(\lambda_m-\lambda_n)>\epsilon_{in}-\epsilon_{im}\right\} \tag{S12}$$

where the approximation follows from a Taylor expansion of $g$ around $\lambda_i$. If we assume that the measurement noise is normally distributed, with mean zero and variance $\sigma^2_{in}$,

$$P^i_{mn} \approx \Phi\left(\frac{g'(\lambda_i)(\lambda_m-\lambda_n)}{\sqrt{\sigma^2_{in}+\sigma^2_{im}}}\right) \tag{S13}$$

and $\Phi(\cdot)$ is the cumulative distribution function of a standard normal random variable. From Eq. S12 and S13, we discern that for small $\lambda_m$ and $\lambda_n$, the probability of correct ordering increases with increasing $g'(\lambda_i)$ and decreasing $\sigma_{im}$ and $\sigma_{in}$. When $g'(\lambda_i) \ll \sqrt{\sigma^2_{in}+\sigma^2_{im}} \approx 0$, $P^i_{m,n}\approx 1/2$. In other words, the *shape* of $g$ in the neighborhood of $\lambda_i$ relative to the scale of measurement noise (relative to the difference in effect sizes) will determine the probability of correct ordering.

##### S2.6.3 An expression for Kendall's tau.

We can use Eq. S12 to find an expression for the expectation of $\hat{\tau}_{ij}$. Letting $\mathbb{1}^k_{mn} := \mathbb{1}\{\hat{Y}_{km}>\hat{Y}_{kn}\}$ (for $k=i,j$),

$$\begin{aligned}
\mathbb{E}[\hat{\tau}_{ij}] &= \frac{2}{|\mathcal{A}_{ij}|(|\mathcal{A}_{ij}|-1)} \sum_{m<n} \mathbb{E}[\mathrm{sgn}(\hat{Y}_{im}-\hat{Y}_{in})\mathrm{sgn}(\hat{Y}_{jm}-\hat{Y}_{jn})] \\
&= \frac{2}{|\mathcal{A}_{ij}|(|\mathcal{A}_{ij}|-1)} \sum_{m<n} \left(1\cdot\mathbb{E}[\mathbb{1}^i_{mn}]-1\cdot(1-\mathbb{E}[\mathbb{1}^i_{mn}])\right)\left(1\cdot\mathbb{E}[\mathbb{1}^j_{mn}]-1\cdot(1-\mathbb{E}[\mathbb{1}^j_{mn}])\right) \\
&= \frac{2}{|\mathcal{A}_{ij}|(|\mathcal{A}_{ij}|-1)} \sum_{m<n} \left(2P^i_{mn}-1\right)\left(2P^j_{mn}-1\right),
\end{aligned} \tag{S14}$$

where $P^i_{mn}$ is defined in Eq. S12. When either $P^i_{mn}$ or $P^j_{mn}=1/2$, the summand is equal to zero.

Let $\mathcal{P}_i$ and $\mathcal{P}_j$ be the sets of pairs of mutations $(m, n)$ for which $P^i_{mn}$ and $P^j_{mn} \approx 1/2$, respectively. This implies that,

$$\mathbb{E}[\hat{\tau}_{ij}] \leq \frac{|\mathcal{P}_i^{\mathsf{c}} \cap \mathcal{P}_j^{\mathsf{c}}|(|\mathcal{P}_i^{\mathsf{c}} \cap \mathcal{P}_j^{\mathsf{c}}| - 1)}{|\mathcal{A}_{ij}|(|\mathcal{A}_{ij}| - 1)}, \tag{S15}$$

where the right hand side is found by assuming that all pairs of mutants in $\mathcal{P}_i^{\mathsf{c}} \cap \mathcal{P}_j^{\mathsf{c}}$ are correctly ordered.

If we further assume that $g$ is monotonically increasing with saturation, i.e., $g'/\sigma_e \approx 0$, *only* at the lower end of the measurement range, then $\lambda_1 \leq \lambda_2 \leq \ldots \leq \lambda_M$ implies that $\mathcal{P}_1 \supseteq \mathcal{P}_2 \supseteq \ldots \supseteq \mathcal{P}_M$. In this regime, Eq. S15 further implies that,

$$\mathbb{E}[\hat{\tau}_{ij}] \leq \frac{\min\left[|\mathcal{P}_i^{\mathsf{c}}|(|\mathcal{P}_i^{\mathsf{c}}| - 1), |\mathcal{P}_j^{\mathsf{c}}|(|\mathcal{P}_j^{\mathsf{c}}| - 1)\right]}{|\mathcal{A}_{ij}|(|\mathcal{A}_{ij}| - 1)}, \tag{S16}$$

and,

$$\max_j \mathbb{E}[\hat{\tau}_{ij}] \leq \frac{|\mathcal{P}_i^{\mathsf{c}}|(|\mathcal{P}_i^{\mathsf{c}}| - 1)}{|\mathcal{A}|(|\mathcal{A}| - 1)}, \tag{S17}$$

where the maximum is taken over all possible mutations and $\mathcal{A}_{ij} = \mathcal{A}$ for all $j$, i.e., there is no missing data.

###### S2.6.4 Approximations to Spearman's correlation under a simplified model

We derive approximate expressions for the maximum and mean Spearman's correlation of each background mutant $i$, under the simplified assumptions detailed at the conclusion of the previous section Section S2.6.3: $g$ is a monotonically increasing function with saturation only at the lower end of the measurement range.

Consider the Spearman's correlation between two backgrounds $i$ and $j$ in which $M = |\mathcal{A}_{ij}|$ mutations are observed,

$$\hat{\rho}_{ij} := \frac{1}{M\hat{\sigma}_i\hat{\sigma}_j} \sum_{m \neq i,j} (\hat{R}_{im} - \bar{R}_{i,\cdot})(\hat{R}_{jm} - \bar{R}_{j,\cdot}), \tag{S18}$$

where $\bar{R}_{i,\cdot}$ and $\bar{R}_{j,\cdot}$ are the average ranks of all mutations in a backgrounds $i$ and $j$, and $\hat{\sigma}_i$ and $\hat{\sigma}_j$ are the standard deviations of the observed ranks, respectively.

When there are no ties or missing measurements, $\bar{R}_{j,\cdot} = \bar{R}_{i,\cdot} = (M+1)/2$ and the variances of the ranks, $\hat{\sigma}_i^2 = \hat{\sigma}_j^2 = (M^2-1)/12$ are likewise equal for all $i$ and $j$, and depend only on $M$ (where the minimum rank is one for convention). Under these assumptions, the Spearman's correlation has a simplified form,

$$\hat{\rho}_{ij} = 1 - \frac{6\sum_{m=1}^{M}[\hat{R}_{im} - \hat{R}_{jm}]^2}{M(M^2-1)}, \tag{S19}$$

that will facilitate calculation of its expected value:

$$\begin{aligned}\mathbb{E}[\hat{\rho}_{ij}] &= 1 - \frac{6}{M(M^2-1)}\left(\sum_{m=1}^{M} \mathbb{E}[\hat{R}_{im}^2] - 2\mathbb{E}[\hat{R}_{im}\hat{R}_{jm}] + \mathbb{E}[\hat{R}_{jm}^2]\right)\\ &= 1 - \frac{12}{M(M^2-1)}\left(\sum_{m=1}^{M} m^2 - \mathbb{E}[\hat{R}_{im}\hat{R}_{jm}]\right),\end{aligned} \tag{S20}$$

and we can solve for the remaining random quantity under some simplifying assumptions.

We refer to the "well-ordered" set of mutations in background $i$ as $\mathcal{W}_i$, where $m \in \mathcal{W}_i$ if $\mathbb{E}[\hat{R}_{im}] = R_m = m$. A site $m$ will be well-ordered when $g'(\lambda_i + \lambda_m)/\sigma_\epsilon \gg 0$. In addition, without loss of generality, we assume that $\lambda_i \leq \lambda_j$, and thus, $\mathcal{W}_i \subseteq \mathcal{W}_j$. Let $\mathbb{1}_{im} := \mathbb{1}\{m \in \mathcal{W}_i\}$. We can then decompose the expectation of the product of double mutant ranks as follows,

$$\begin{aligned}&\sum_{m=1}^{M} \mathbb{E}[\hat{R}_{im}\hat{R}_{jm}]\\ &\quad= \sum_{m=1}^{M} m^2 \mathbb{1}_{im}\mathbb{1}_{jm} + m\left(\frac{|\mathcal{W}_i^{\mathsf{c}}|+1}{2}\right)\mathbb{1}_{jm}(1-\mathbb{1}_{im}) + \left(\frac{|\mathcal{W}_j^{\mathsf{c}}|+1}{2}\right)\left(\frac{|\mathcal{W}_i^{\mathsf{c}}|+1}{2}\right)(1-\mathbb{1}_{jm})(1-\mathbb{1}_{im})\\ &\quad= \sum_{m=|\mathcal{W}_i^{\mathsf{c}}|+1}^{M} m^2 + \left(\frac{|\mathcal{W}_i^{\mathsf{c}}|+1}{2}\right)\sum_{m=|\mathcal{W}_j^{\mathsf{c}}|+1}^{|\mathcal{W}_i^{\mathsf{c}}|} m + \sum_{m=1}^{|\mathcal{W}_j^{\mathsf{c}}|}\left(\frac{|\mathcal{W}_i^{\mathsf{c}}|+1}{2}\right)\left(\frac{|\mathcal{W}_j^{\mathsf{c}}|+1}{2}\right)\\ &\quad= \frac{M(M+1)(2M+1)}{6} - \frac{1}{12}|\mathcal{W}_i^{\mathsf{c}}|(|\mathcal{W}_i^{\mathsf{c}}|^2 - 1),\end{aligned} \tag{S21}$$

and thus,

$$\mathbb{E}[\hat{\rho}_{ij}] = 1 - \frac{|\mathcal{W}_i^{\mathsf{c}}|(|\mathcal{W}_i^{\mathsf{c}}|^2 - 1)}{M(M^2-1)}, \tag{S22}$$

and as with Kendall's $\tau$, the expected value of $\hat{\rho}_{ij}$ depends only on the minimum size of the well-ordered sets, $\mathcal{W}_i$ and $\mathcal{W}_j$.

###### S2.6.5 Average rank in a simplified model

We can similarly find an expression for the average rank of a mutation $m$ across all backgrounds $i$,

$$\mathbb{E}[\bar{R}_{\cdot m}] \coloneqq \frac{1}{M}\sum_{i\neq m}\mathbb{E}[\hat{R}_{im}] = \frac{1}{M}\left[\sum_{\substack{i \text{ s.t.}\\ m\in\mathcal{W}_i}} m + \sum_{\substack{i \text{ s.t.}\\ m\notin\mathcal{W}_i}} \frac{|\mathcal{W}_i^{\mathsf{c}}|+1}{2}\right]. \tag{S23}$$

Letting $W_m \coloneqq |\{i \text{ s.t. } m \notin \mathcal{W}_i\}|$,

$$\mathbb{E}[\bar{R}_{\cdot m}] = \frac{1}{M}\left[\sum_{i=W_m+1}^{M} m + \sum_{i=1}^{W_m}\frac{|\mathcal{W}_i^{\mathsf{c}}|+1}{2}\right] = \left(\frac{M-W_m}{M}\right)m + \frac{1}{2M}\left(\frac{W_m(W_m+1)}{2} + \sum_{i=1}^{W_m}|\mathcal{W}_i^{\mathsf{c}}|\right), \tag{S24}$$

where, clearly, when $W_m = 0$, $\mathbb{E}[\bar{R}_m] = m$.

The above calculation codifies the intuition that, under the assumption of GE, $\mathbb{E}[\bar{R}_{\cdot m}]$ will be an increasing function of the single mutant ranks $\hat{R}_m$. ($\lambda_m \leq \lambda_{m+1} \implies W_m \geq W_{m+1} \implies \mathbb{E}[\bar{R}_{\cdot,m+1}] \geq \mathbb{E}[\bar{R}_{\cdot m}]$.) This conclusion motivates the definition of a mutants *residual* as the difference between the ranked average mutant effect, $R[\bar{R}_{\cdot m}]$, and its single effect $\hat{R}_m$.

###### S2.6.6 Effect of specific epistasis on ordering

We derive conditions under which SE will result in deviations from the null model of single-trait GE. In the presence of SE, the estimated phenotype of double mutant $im$ is given by,

$$\hat{Y}_{im} = g(\lambda_i + \lambda_m + \lambda_{im}) + \epsilon_{im}, \tag{S25}$$

where $\lambda_{im} \neq 0$ when the two mutations are specifically epistatic. To understand the effect of SE on the rank statistics of double mutations, we again consider pairwise ordering of two mutations $m$ and $n$ in the background of mutation $i$. In the limit where $\lambda_m + \lambda_{im}$ and $\lambda_n + \lambda_{in}$ are small,

$$\begin{aligned} P_{m,n}^{(i)} \coloneqq \mathbb{P}\{\hat{Y}_{im} > \hat{Y}_{in}\} &= \mathbb{P}\{g(\lambda_i + \lambda_m + \lambda_{im}) - g(\lambda_i + \lambda_n + \lambda_{in}) > \epsilon_{in} - \epsilon_{im}\} \\ &\approx 1 - \Phi\left(\frac{\tilde{g}'(\lambda_i)(\lambda_m - \lambda_n + \lambda_{im} - \lambda_{in})}{\sigma_\epsilon}\right) \end{aligned} \tag{S26}$$

where, for exposition, $\epsilon_{in} - \epsilon_{im} \sim \mathcal{N}(0, \sigma_\epsilon^2)$. When epistasis is sparse, we can assume, without loss of generality, that $\lambda_{in} = 0$, and thus, when $g'(\lambda_i)/\sigma_\epsilon \gg 0$, $P_{m,n}^{(i)} \geq 1/2$ if $\lambda_{im} > \lambda_m - \lambda_n$. Below, we relate the observed rank deviation due to SE to the magnitude and direction of the underlying epistatic effect $\lambda_{im}$.

###### S2.6.7 Relating rank deviations to estimates of the epistatic effect

We can use the intuition developed in Section S2.6.6 to approximate the magnitude and direction of a given interaction term $\lambda_{ij} > 0$. Consider a mutant pair with single and double mutant ranks, $\hat{R}_i$, $\hat{R}_m$, $\hat{R}_{im}$, and $\hat{R}_{mi}$. A rank of $\hat{R}_{im} = R_*$ implies,

$$\{\hat{Y}_{im} > \hat{Y}_{in}\} \ \forall\ m \text{ s.t. } \hat{R}_{in} < R_\star, \tag{S27}$$

which further implies,

$$\{g(\lambda_i + \lambda_m + \lambda_{im}) - g(\lambda_i + \lambda_n) > \epsilon_{in} - \epsilon_{im}\} \approx \{g'(\lambda_i + \lambda_n)[\lambda_{im} - (\lambda_n - \lambda_m)] > \epsilon_{in} - \epsilon_{im}\}, \tag{S28}$$

where the approximation is valid for small $\lambda_m + \lambda_{im} - \lambda_n$, and we let $\lambda_{in} = 0$ under the assumption of sparsity. Letting $n^* \coloneqq \arg\max_n \hat{R}_{in} < R_*$,

$$\{\hat{Y}_{im} > \hat{Y}_{in^*}\} \implies \left\{\lambda_{im} > \frac{\epsilon_{in^*} - \epsilon_{im}}{g'(\lambda_i + \lambda_{n^*})} + (\lambda_{n^*} - \lambda_m)\right\}, \tag{S29}$$

which, for large $g'(\lambda_i + \lambda_{n^*})$, Eq. S29 yields an intuitive result: For $\hat{R}_{im}$ to exceed $R_{in^*}$, $\lambda_{im}$ must be greater than the difference of the first order effects of mutants $n^*$ and $m$ on fitness.

We can derive an analogous result for $\lambda_{im} < 0$. Letting $n^* \coloneqq \arg\min_n \hat{R}_{in} > R_*$,

$$\left\{\lambda_{im} < \frac{\epsilon_{in*} - \epsilon_{im}}{g'(\lambda_i + \lambda_{n^*})} + (\lambda_{n^*} - \lambda_m)\right\}, \tag{S30}$$

and likewise, for large $g'(\gamma_i + \gamma_n^*)$, the epistatic effect must be smaller than the difference between the single effects, $\lambda_{n^*} - \lambda_m$.

Eq. S29 and S30 further suggest how detection of SE depends on the shape of the nonlinearity relative to the measurement noise. When the slope of the nonlinearity approaches zero, the quantity on the right hand sides of Eq. S29 and S30 blows up—deviations in rank due to noise are large—and the typical magnitude of $\lambda_{im}$ must likewise be larger to achieve statistical significance.

##### S2.6.8 Asymmetry in statistical power

Application of R&R to simulated and empirical data suggest that power to detect positive versus negative interactions is asymmetric in a large portion of the measurement range. While limited power to detect positive epistasis among the most beneficial mutant pairs and negative epistasis among the most deleterious mutant pairs is readily explained (see Section 3.1), an explanation for the bias toward negative epistasis among pairs of beneficial and deleterious mutants, as observed in the analysis of the GB1-like simulations and the empirical datasets, is less immediate (see Figs. 2 and 3).

Consider two mutations $i$ and $j$ with epistatic interaction of magnitude $\lambda_{ij}$. Asymmetry in the power to detect SE would occur if,

$$\mathbb{E}\left[|\hat{D}_{ij}||\lambda_i, \lambda_j, -\lambda_{ij}\right] > \mathbb{E}\left[|\hat{D}_{ij}||\lambda_i, \lambda_j, \lambda_{ij}\right], \tag{S31}$$

or vice versa. Though, we consider only the case where Eq. S31 holds. If we assume that the single mutants are well-ordered, we can express Eq. S31 as,

$$\mathbb{E}\left[|\hat{R}_{ij} + \hat{R}_{ji} - R_i - R_j||\lambda_i, \lambda_j, -\lambda_{ij}\right] > \mathbb{E}\left[|\hat{R}_{ij} + \hat{R}_{ji} - R_i - R_j||\lambda_i, \lambda_j, \lambda_{ij}\right].$$

If the probability of observing an increase in double mutant rank in the presence of negative SE ($-\lambda_{ij} < 0$) and a decrease in double mutant rank for positive SE ($\lambda_{ij} > 0$) are negligible, we have,

$$-\mathbb{E}\left[\hat{R}_{ij} + \hat{R}_{ji}|\lambda_i, \lambda_j, -\lambda_{ij}\right] + R_i + R_j > \mathbb{E}\left[\hat{R}_{ij} + \hat{R}_{ji}|\lambda_i, \lambda_j, \lambda_{ij}\right] - R_i - R_j. \tag{S32}$$

Let the lower detection limit of the assay be $\Lambda^*$, where $\Lambda^* \coloneqq \arg\max_\Lambda \left\{\mathbb{P}\{\hat{N}^1 = 0|\Lambda\} \approx 1\right\}$. For the purposes of this analysis, we will not include pseudo-counts in the fitness estimates. Thus, all variants with post-selection reads equal to zero ($\hat{N}_{ij} = 0$) will be mapped to the same rank.

Suppose, $i$ is a very deleterious mutation, i.e., $\lambda_i \leq \Lambda^*$, while $j$ is a neutral or beneficial mutation, i.e., $\lambda_j > \Lambda^*$. Further suppose, $\lambda_i + \lambda_j \geq \Lambda^*$. If, $\Lambda_{ij}^- \coloneqq \lambda_i + \lambda_j - \lambda_{ij} < \Lambda^*$, then $\mathbb{E}[\hat{R}_{ij}|\Lambda_{ij}^-] \approx M_i/2$, where $M_i$ is the total number of variants $k \neq i$ mapped to zero in the background of mutant $i$ (using zero-indexing for the ranks). Similarly, $\mathbb{E}[\hat{R}_{ji}|\Lambda_{ij}^-] \approx M_j/2$.

Under our assumptions, $\Lambda_{ij}^+ \coloneqq \lambda_i + \lambda_j + \lambda_{ij} > \Lambda^*$, and thus,

$$\mathbb{E}[\hat{R}_{ij}|\Lambda_{ij}^+] \approx \sum_{k \neq i,j} \mathbb{1}\{\lambda_{ij} > \lambda_k - \lambda_j\} \approx R_j + \sum_{R_k > R_j} \mathbb{1}\{\lambda_{ij} > \lambda_k - \lambda_j\},$$

where we have assumed, for simplicity, that $\lambda_{kj} = 0$ for all $k \neq i$, and have ignored noise in the fitness measurements. Similarly,

$$\mathbb{E}[\hat{R}_{ji}|\Lambda_{ij}^+] \approx \sum_{k \neq i,j} \mathbb{1}\{\lambda_{ij} > \lambda_k - \lambda_i\} \approx R_i + \sum_{R_k > R_i} \mathbb{1}\{\lambda_{ij} > \lambda_k - \lambda_i\}.$$

Combining these expressions, we can now express the condition in Eq. S31 as,

$$-\left(\frac{M_i}{2} + \frac{M_j}{2}\right) + R_i + R_j > \sum_{R_k > R_j} \mathbb{1}\{\lambda_{ij} > \lambda_k - \lambda_j\} + \sum_{R_k > R_i} \mathbb{1}\{\lambda_{ij} > \lambda_k - \lambda_i\}.$$

If we assume that $R_i \approx 0$ and $R_j \approx M$, where $M$ is the total number of mutants,

$$M - \frac{M_i}{2} \gtrapprox \frac{M_j}{2} + \sum_{R_k > M_j} \mathbb{1}\{\lambda_{ij} > \lambda_k - \lambda_j\} \approx \sum_{R_k > M_j} \mathbb{1}\{\lambda_{ij} > \lambda_k - \lambda_j\}, \tag{S33}$$

where, we have assumed that $R_j \approx M \implies M_j \approx 0$. We observe that the left hand side of Eq. S33 does not depend on the magnitude of $\lambda_{ij}$, only the number of mutants mapped to zero in the background of $i$. On the other hand, the right hand side depends on the magnitude of $\lambda_{ij}$. Unless $\lambda_{ij}$ is large relative to differences between the single effects, the left hand side will be greater (for modest values of $M_i$).
